# Human online adaptation to changes in prior probability

**DOI:** 10.1101/483842

**Authors:** Elyse H. Norton, Luigi Acerbi, Wei Ji Ma, Michael S. Landy

**Affiliations:** Psychology Department, New York University, New York, NY, USA; Center for Neural Science, New York University, New York, NY, USA

**Author notes:** Current Address: Département des neurosciences fondamentales, Université de Genève, CMU, 1 rue Michel-Servet, 1206 Genève, Switzerland.

## Abstract

Optimal sensory decision-making requires the combination of uncertain sensory signals with prior expectations. The effect of prior probability is often described as a shift in the decision criterion. Can observers track sudden changes in probability? To answer this question, we used a change-point detection paradigm that is frequently used to examine behavior in changing environments. In a pair of orientation-categorization tasks, we investigated the effects of changing probabilities on decision-making. In both tasks, category probability was updated using a sample-and-hold procedure. We developed an ideal Bayesian change-point detection model in which the observer marginalizes over both the current run length (i.e., time since last change) and the current category probability. We compared this model to various alternative models that correspond to different strategies – from approximately Bayesian to simple heuristics – that the observers may have adopted to update their beliefs about probabilities. We find that probability is estimated following an exponential averaging model with a bias towards equal priors, consistent with a conservative bias. The mechanism underlying change of decision criterion is a combination of on-line estimation of prior probability and a stable, long-term equal-probability prior, thus operating at two very different timescales.

**Author summary:** We demonstrate how people learn and adapt to changes to the probability of occurrence of one of two categories on decision-making under uncertainty. The study combined psychophysical behavioral tasks with computational modeling. We used two behavioral tasks: a typical forced-choice categorization task as well as one in which the observer specified the decision criterion to use on each trial before the stimulus was displayed. We formulated an ideal Bayesian change-point detection model and compared it to several alternative models. We found that the data are best fit by a model that estimates category probability based on recently observed exemplars with a bias towards equal probability. Our results suggest that the brain takes multiple relevant time scales into account when setting category expectations.

## Introduction

Sensory decision-making involves making decisions under uncertainty. Furthermore, optimal sensory decision-making requires the combination of uncertain sensory signals with prior expectations. Perceptual models of decision-making often incorporate prior expectations to describe human behavior. In Bayesian models, priors are combined with likelihoods to compute a posterior [1]. In signal detection theory, the effect of unequal probabilities (signal present vs. absent) is a shift of the decision criterion [2].

The effects of prior probability on the decision criterion have been observed in detection [2–4], line tilt [5], numerosity estimation [6, 7], recognition memory [8], and perceptual categorization [9] tasks, among others. These studies generally use explicit priors, assume a fixed effect, and treat learning as additional noise. However, there are many everyday tasks in which the probability of a set of alternatives needs to be assessed based on one’s past experience with the outcomes of the task. The importance of experience has been demonstrated in studies examining differences between experience-based and description-based decisions [10, 11] and in perceptual-categorization tasks with unequal probability, in which response feedback leads to performance that is closer to optimal than observational feedback [12, 13]. While these studies demonstrate the importance of experience on decision-making behavior, they do not describe how experience influences expectation formation. The influence of experience on the formation of expectations has been demonstrated for learning stimulus mean [14–17], variance [14, 18], higher-order statistics [19], likelihood distributions [20], the rate of change of the environment [15–17, 21–23], and prior probability [24, 25]. Here, we add to previous work by investigating how one’s previous experience influences probability learning in a changing environment.

In the previous work on probability learning by Behrens et al. [24], participants tracked the probability of a reward to learn the value of two alternatives. This is a classic decision-making task that involves combining prior probability and reward. In contrast, we are interested in perceptual decision-making under uncertainty, in which prior probability is combined with uncertain sensory signals. We might expect differences in strategy between cognitive and perceptual tasks, as cognitive tasks can be susceptible to additional biases. For example, participants often exhibit base rate neglect (i.e., they ignore prior probability when evaluating evidence) in cognitive tasks [26] but not in perceptual tasks [2]. On the other hand, Behrens et al. [24] found that participants’ behavior was well described by an optimal Bayesian model, in that observed learning rates quantitatively matched those of a Bayesian decision-maker carrying out the same task. Although it is important to note that behavior was not compared to alternative models. A more recent study by Zylberberg et al. [25] examined probability learning under uncertainty in a motion-discrimination task. In this study, probability was estimated from a confidence signal rather than explicit feedback. Additionally, probability was changed across blocks, allowing participants to infer a change had occurred. Here, we examine probability learning when feedback is explicit and changes are sudden.

To investigate probability learning in uncertain and changing conditions, we designed a perceptual-categorization task in which observers need to learn category probability through experience. To prevent the use of external cues (e.g., the start of a new block indicating a change in probability) probabilities were changed using a sample-and-hold procedure. This approach has been used in decision-making [21, 22, 24] and motor domains [18] to examine behavior in dynamic contexts. Observers completed both a covert- and overt-criterion task, in which the decision criterion was implicit or explicit, respectively. The overt task, previously developed by Norton et al. [16], provided a richer dataset upon which to compare models of human behavior. We determined how observers tracked category probability in a changing environment by comparing human behavior to both Bayesian and alternative models. We find that a number of models qualitatively describe human behavior, but that, quantitatively, all models are outperformed by an exponential averaging model with a bias towards equal priors. Our results suggest that changes in the decision criterion are the result of both probability estimates computed on-line and a more stable, long-term prior.

## Results

### Experiment

During each session, observers completed one of two orientation-categorization tasks. On each trial in the *covert-criterion* task, observers categorized an ellipse as belonging to category A or B by key press (Fig 1A). On each trial in the *overt-criterion* task, observers used the mouse to rotate a line to indicate their criterion prior to the presentation of an ellipse (Fig 1B). The observer was correct if the ellipse belonged to category A and was clockwise of the criterion line or if the ellipse belonged to category B and was counter-clockwise of the criterion line. The overt-criterion task is an explicit version of the covert-criterion task developed by Norton et al. [16]. The overt-criterion task provides a richer dataset than the covert-criterion task in that it affords a continuous measure and allows us to see trial by trial changes in the reported decision criterion, at the expense of being a more cognitive task. In both tasks, the categories were defined by normal distributions on orientation with different means (*µ*_B_ = *µ*_*A*_ + Δ*θ*) and equal standard deviation (*σ*_*C*_); the mean of category A was set clockwise of the the mean of category B and a neutral criterion would be located halfway between the category means (Fig 1C). The difficulty of the task was equated across participants by setting Δ*θ* to a value that predicted a *d*^′^ value of 1.5 based on the data from the initial measurement session (see Methods). All data are reported relative to the neutral criterion, *z*_neutral_ = (*µ*_A_ + *µ*_B_)/2.

**Fig 1.**
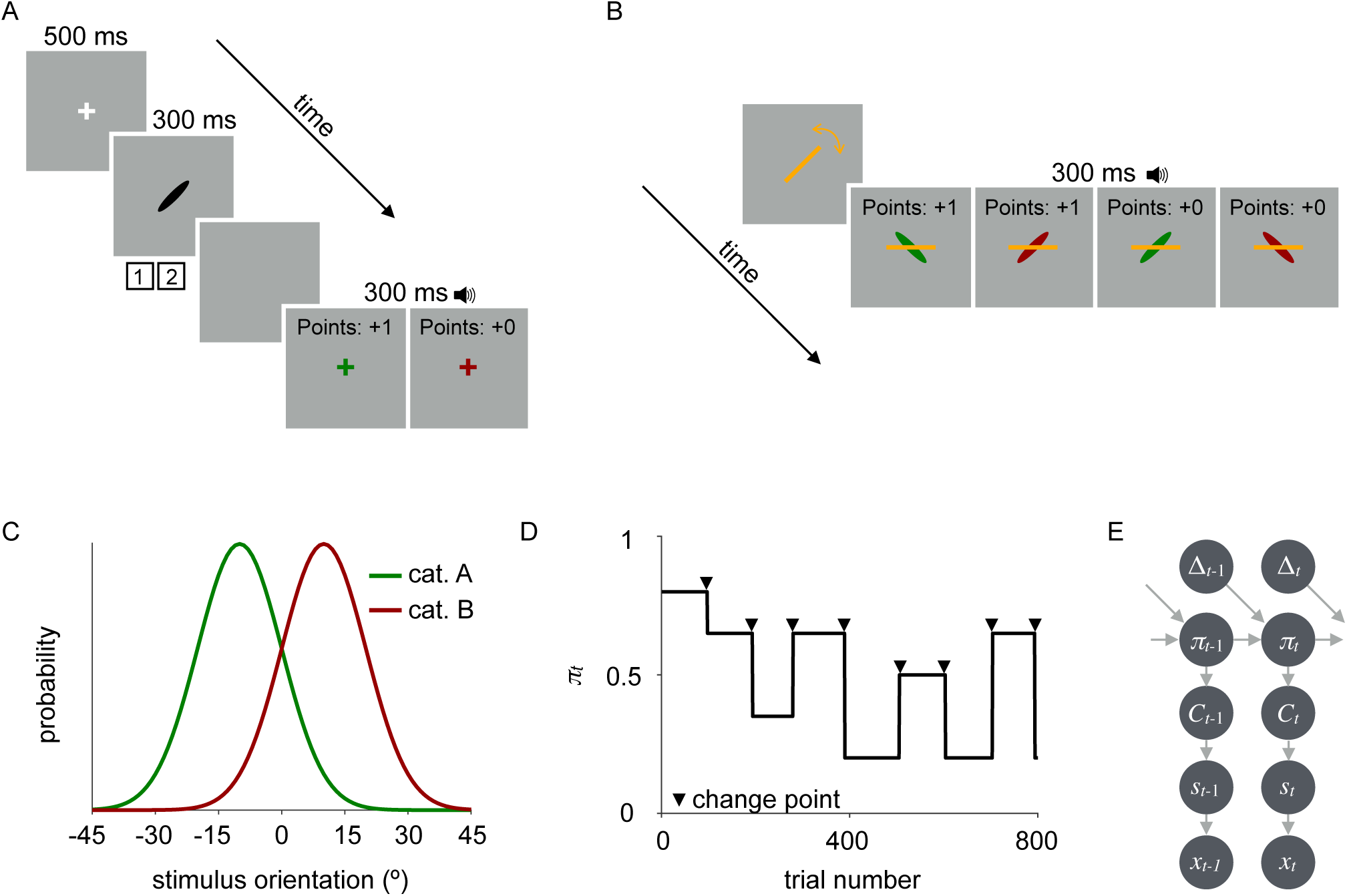
Experimental design. A: Trial sequence in the covert-criterion task. After stimulus offset, observers reported the category by key press and received auditory feedback indicating the correctness of their response. In addition, the fixation cross was displayed in the color of the generating category. B: Trial sequence in the overt-criterion task. Prior to stimulus onset, observers rotated a line to indicate their criterion by sliding the mouse side-to-side. When the observer clicked the mouse, a stimulus was displayed under the criterion line and auditory feedback was provided. C: Example stimulus distributions for category A (green) and category B (red). Note that numbers on the *x*-axis are relative to the neutral criterion (i.e., 0 is the neutral criterion). D: Example of random stepwise changes in probability across a block of trials. Change points occurred every 80-120 trials and are depicted above by the black arrows. Category A probabilities *π*_*t*_ were sampled from a discrete set, *S*_*π*_ = {0.2, 0.35, 0.5, 0.65, 0.8}. E: Generative model for the task in which category probability *π* is updated following a sample-and-hold procedure, a category *C* is selected based on the category probability, a stimulus *s* is drawn from the category distribution and is corrupted by visual noise resulting in the noisy measurement *x*. Note that this diagram omits the dependency that leads to change points every 80-120 trials.

Prior to testing, observers were trained on the categories. Only the covert-criterion task was used for training (see Section 6 in S1 Appendix). During training, category probability was equal (*π* = 0.5) and observers received feedback on every trial that indicated both the correctness of the response (tone) and the generating category (visual). As a measure of category learning, we compute the probability of being correct in the training block and averaged across sessions. All observers learned the categories to the expected level of accuracy (*p*(correct) = 0.74 ± 0.01; mean ± SEM across observers), given that the expected fraction of correct responses for an ideal observer with *d*^′^ = 1.5 and equal priors over categories is 0.77. As an additional test of category learning, immediately following training observers estimated the mean orientation of each category by rotating an ellipse. Each category mean was estimated once and no feedback was provided. There was a significant correlation between the true and estimated means for each category (category A: *r* = 0.82, *p <* 0.0001; category B: *r* = 0.97, *p <* 0.0001), suggesting that categories were learned. However, on average mean estimates were repelled from the category boundary (average category A error of 11.3° ± 6.3° and average category B error of −8.0° ± 2.6°; mean ± SEM across observers) suggesting a systematic repulsive bias.

To determine how category probability affects decision-making, during testing category A probability *π*_*t*_ was determined using a sample-and-hold procedure (Fig 1D; category B probability was 1 − *π*_*t*_). For *t* = 1, category A probability was randomly chosen from a set of five probabilities *S*_*π*_ = {0.2, 0.35, 0.5, 0.65, 0.8}. On most trials, no change occurred (Δ_*t*_ = 0), so that *π*_*t*+1_ = *π*_*t*_. Every 80-120 trials there was a change point (Δ_*t*_ = 1), with change point sampled uniformly. At each change point, category probability was randomly selected from the *S*_*π*_ excluding the current probability. On each trial *t*, a category *C*_*t*_ was randomly selected (with *P* (category A) = *π*_*t*_) and a stimulus *s*_*t*_ was drawn from the stimulus distribution corresponding to the selected category. We assume that the observer’s internal representation of the stimulus is a noisy measurement *x*_*t*_ drawn from a Gaussian distribution with mean *s*_*t*_ and standard deviation *σ*_v_, which represents visual noise (v). The generative model of the task is summarized in Fig 1E.

### Bayesian models

To understand how decision-making behavior is affected by changes in category probability, we compared observer performance to several Bayesian models. To compute the behavior of a Bayesian observer, we developed a Bayesian change point-detection algorithm, based on Adams and MacKay [27], but which also accounts for Markov dependencies in the transition distribution after a change. Specifically, the Bayesian observer estimates the posterior distribution over the current run length (time since the last change point), and the state (category probability) before the last change point, given the data so far observed (category labels until trial *t*, C_*t*_ = (*C*_1_*, …, C*_*t*_)). We denote the current run length at the end of trial *t* by *r*_*t*_, the current state by *π*_*t*_, and the state before the last change point by *ξ*_*t*_, where both *π*_*t*_*, ξ*_*t*_ ∈ *S*_*π*_. That is, if a changepoint occurs after trial *t* (i.e., *r*_*t*_ = 0), then the new category A probability will be *π*_*t*_ and the previous run’s category probability *ξ*_*t*_ = *π*_*t* − 1_. If no changepoint occurs, both *π* and *ξ* remain unchanged. We use the notation 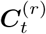 to indicate the set of observations (category labels) associated with the run *r*_*t*_, which is 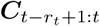 for *r*_*t*_ > 0, and ∅ for *r*_*t*_ = 0. The range of times with a colon, e.g., 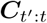, indicates the sub-vector of C with elements from *t*^′^ to *t* included.

Both of our tasks provide category feedback, so that at the end of trial *t* the observer has been informed of *C*_1:*t*_. In S1 Appendix we derive the iterative Bayesian ideal-observer model. After each trial, the model calculates a posterior distribution over possible run lengths and previous probability states, *P* (*r*_*t*_*, ξ*_*t*_|*C*_1:*t*_). The generative model makes it easy to calculate the conditional probability of the current state for a given run length and previous state, *P* (*π*_*t*_|*r*_*t*_*, ξ*_*t*_, *C*_1:*t*_). These two distributions may be combined, marginalizing (summing) across the unknown run length and previous states to yield the predictive probability distribution of the current state, *P* (*π*_*t*_|*C*_1:*t*_). Given this distribution over states, in both tasks the observer needs to determine the probability of each category. In particular,

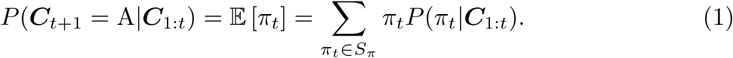

In the overt task, the ideal observer sets the current criterion to the optimal value 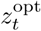 based on the known category orientation distributions and the current estimate of category probabilities. Further, in the ideal and all subsequent models of the overt task, in addition to early sensory noise (*σ*_v_) we assume the actual setting is perturbed by adjustment noise (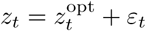 where 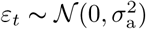).

In the covert task, the observer views a stimulus and makes a noisy measurement *x*_*t*_ of its true orientation *s*_*t*_ with noise variance 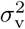. The prior category probability is combined with the noisy measurement to compute category A’s posterior probability *P* (*C*_*t*+1_ = A|*x*_*t*+1_, *C*_1:*t*_). The observer responds “A” if that probability is greater than 0.5.

We consider the ideal-observer model (Bayes_ideal_) and three (suboptimal) variants thereof, which deviate from the ideal observer in terms of their beliefs about specific features of the experiment (Bayes_*r*_, Bayes_*π*_, and Bayes_*β*_). Two further variants of the Bayesian model (Bayes_bias_ and Bayes_*r,β*_) are described in S1 Appendix. Crucially, all these models are “Bayesian” in that they compute a posterior over run length and probability state, but they differ with respect to the observer’s assumptions about the generative model.

#### Bayes_ideal_

The ideal Bayesian observer uses knowledge of the generative model to maximize expected gain. This model assumes knowledge of sensory noise, the category distributions, the set of potential probability states, and the run length distribution. Because observers were trained on the categories prior to completing each categorization task, assuming knowledge of the category distributions seems reasonable. Further, it is reasonable to assume knowledge of sensory noise as this is a characteristic of the observer. However, since observers were not told how often probability would change, nor were they told the set of potential probability states, observers may have had incorrect beliefs about the generative model. Thus, we developed additional Bayesian models (described below), in which observers could have potentially incorrect beliefs about different aspects of the generative model.

#### Bayes_*r*_

The Bayes_*r*_ model allows for an observer to have an incorrect belief about the run length distribution. For a given discrete *r*, the observer believes that the run length distribution is drawn from a discrete uniform distribution, ∼ Unif 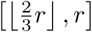. We chose this particular interval rather than a more general one, because it is simple and includes the true generative distribution. For the ideal observer, *r* = 120.

#### Bayes_*π*_

The Bayes_*π*_ model allows for an observer to have an incorrect belief about the set of probability states. The veridical set of experimental states is five values of *π* ranging from 0.2 to 0.8. The Bayes_*π*_ model observer also assumes five possible values of *π* evenly spaced, but ranging from *π*_min_ to *π*_max_ = 1 − *π*_min_, where *π*_min_ is a free parameter.

#### Bayes_*β*_

While incorrect assumptions about parameters of the generative model result in suboptimal behavior, suboptimality can also arise from prior biases (that is, incorrect hyper-parameters). The Bayes_*β*_ model, like the Bayes_ideal_ model, assumes knowledge of the generative model, but also includes a hyperprior Beta(*β, β*) that is applied after a change-point. Thus, as *β* increases, the posterior belief over category probabilities is biased toward equal probability. For the ideal observer, *β* = 1 (a uniform distribution).

### Heuristic models

In addition to the Bayesian models described above, we tested the following heuristic models that do not require the observer to compute a posterior over run length and probability state. In each of the following models, assumptions vary about whether and how probability is estimated. In the Fixed Criterion (Fixed) model the observer assumes fixed probabilities. In the Exponential-Averaging (Exp), Exponential-Averaging with Prior Bias (Exp_bias_), and the Wilson et al. (2013) models, probability is estimated based on the recent history of categories. Finally, in the Reinforcement-Learning (RL) model, the decision criterion is updated following an error-driven learning rule with no assumptions about probability. We also tested three additional heuristic models that are described in the Supplementary material (see S1 Appendix).

Each of the following models, other than the RL model, computes an estimate of category probability 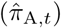 on each trial and the estimated probability of the alternative is 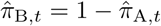. On each trial, the optimal criterion *z*_opt_ is computed based on these estimated probabilities in the identical manner as for the Bayesian models. To make a categorization decision in the covert-criterion task, the criterion is applied to the noisy observation of the stimulus. In the overt-criterion task, the observer reports the criterion, which we again assume is corrupted by adjustment noise.

#### Fixed

While incorporating category probability into the decision process maximizes expected gain, an alternative strategy is to ignore changes in probability and fix the decision criterion throughout the block. In the fixed-criterion model, we assume equal category probability and the criterion is set to the neutral criterion:

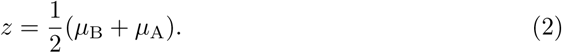

This model is equivalent to a model in which the likelihood ratio, 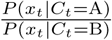, is used to make categorization decisions rather than the posterior odds ratio in the covert-criterion task. This is a reasonable strategy for an observer who wants to make an informed decision, but is unsure about the current probability state and its rate of change.

#### Exp

The exponential-averaging model computes smoothed estimates of category probability by taking a weighted average of previously experienced category labels, giving more weight to recently experienced labels:

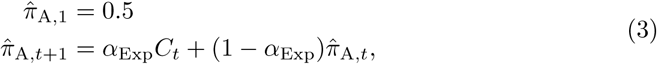

where *α*_Exp_ is the smoothing factor, 0 *< α*_Exp_ < 1, and *C*_*t*_ = 1 if category A is observed on trial *t* and *C*_*t*_ = 0 otherwise. The time constant of memory decay for this model is 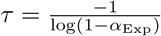 Mathematically this model is equivalent to a delta-rule [28] model based on an “error” that is the difference between the current category and the current probability estimate: 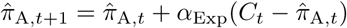 The criterion *z* is then set to the optimal value based on this category-probability estimate.

#### Exp_bias_

The Exp_bias_ model is identical to the Exp model, while also incorporating a common finding in the literature [2] known as conservatism (i.e., observers are biased towards the neutral criterion). On each trial an estimate of probability is computed as described in (Eq 3), and averaged with a prior probability of 0.5:

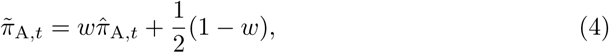

where *w* is the weight given to the probability estimate 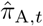 and (1 − *w*) is the weight given to *π*_A_ = 0.5. The criterion *z* is also set to the optimal value based on this category-probability estimate.

#### Wilson et al. (2013)

Due to the complexity of the full Bayesian change-point detection model, Wilson et al. [29] developed an approximation to the full model using a mixture of delta rules. In their reduced model, the full run-length distribution is approximated by maintaining a subset of all possible run lengths. These are referred to as nodes {*l*_1_*, …, l*_*i*_}. On each trial, an estimate of probability is computed for each node,

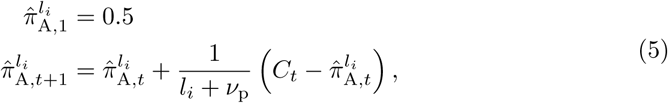

where *ν*_p_ is a model parameter that represents the strength of the observer’s prior towards equal category probability (pseudocounts of a Beta prior; larger *V*_p_ corresponds to stronger conservatism) and *C*_*t*_ is the category label. The learning rate for each node is thus *α* wilson, 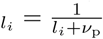. For a single-node model, this is identical to the Exp model described above (Eq 3). To obtain an overall estimate of probability in a multiple-node model, estimates (Eq 5) are combined by taking a weighted average. The weights, *p*(*l*_*i*_|*C*_1:*t*_), are updated on every trial. The update is dependent on an approximation to the change-point prior, which in turn depends on the hazard rate *h* (for details see Wilson et al. [29], Eqs 25-31, along with the later correction [30]). In other words, when the probability of a change is high, more weight is given to *l*_1_, which results in a greater change in the overall probability estimate. But when the probability of a change is low, more weight is given to nodes greater than *l*_1_ and the probability estimate is more stable. For a three-node model probability is estimated as

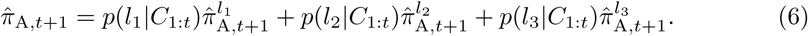

For the purpose of the current experiment, we used a three-node model in which the first node was fixed (*l*_1_ = 1) and *l*_2_ and *l*_3_ were allowed to vary. In addition, the prior strength parameter, *ν*_p_, which modulates the learning rate, was also free. By allowing *ν*_p_ to vary, this model is equivalent to the three-node model described by Wilson and colleagues [29] in which all nodes were free and *ν*_p_ = 2 was fixed. The hazard rate was set to 0.01 and we assumed a change occurred at *t* = 1, so that all the weight was given to *l*_1_. We also tested an alternative model with a fixed value of *V*_p_ = 2, but it resulted in a worse fit.

#### RL

While the models described above make assumptions about how probability is estimated, it is also possible that observers simply update the decision criterion without estimating the current probability state. Reinforcement-learning is a model-free approach that assumes the observer does not use knowledge of the environmental structure. Instead, the decision criterion, rather than an estimate of category probability, is updated. The observer updates the internal criterion (*z*) on each trial according to the following delta rule:

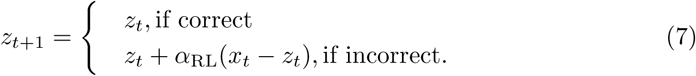

Thus, the criterion is updated when negative feedback is received by taking a small step in the direction of the difference between the noisy observation (*x*_*t*_) and current criterion (*z*_*t*_), where the step size is scaled by the learning rate *α*_RL_. Nothing is changed after a correct response. Assuming effective training, we initialize the RL model by setting 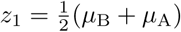.

### Raw data

Fig 2 shows raw data for single observers in the covert (Fig 2A) and overt (Fig 2C) tasks. For visualization in the covert task, the ‘excess’ number of A responses is plotted as a function of trial (gray line in Fig 2A). To compute the ‘excess’ number of A responses, we subtracted 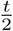 from the cumulative number of A responses. Thus, ‘excess’ A responses are constant for an observer who reported A and B equally often, increase when A is reported more, and decrease when A is reported less. To get a sense of how well the observer performed in the covert task, the number of ‘excess’ A trials (based on the actual category on each trial rather than the observer’s response) is shown in black (Fig 2A, top). For reference, *π*_A_ is shown as a function of trial (Fig 2A, bottom). From visual inspection, the observer reported A more often when *π*_A, *t*_ > 0.5 and B more often when *π*_A, *t*_ < 0.5.

**Fig 2.**
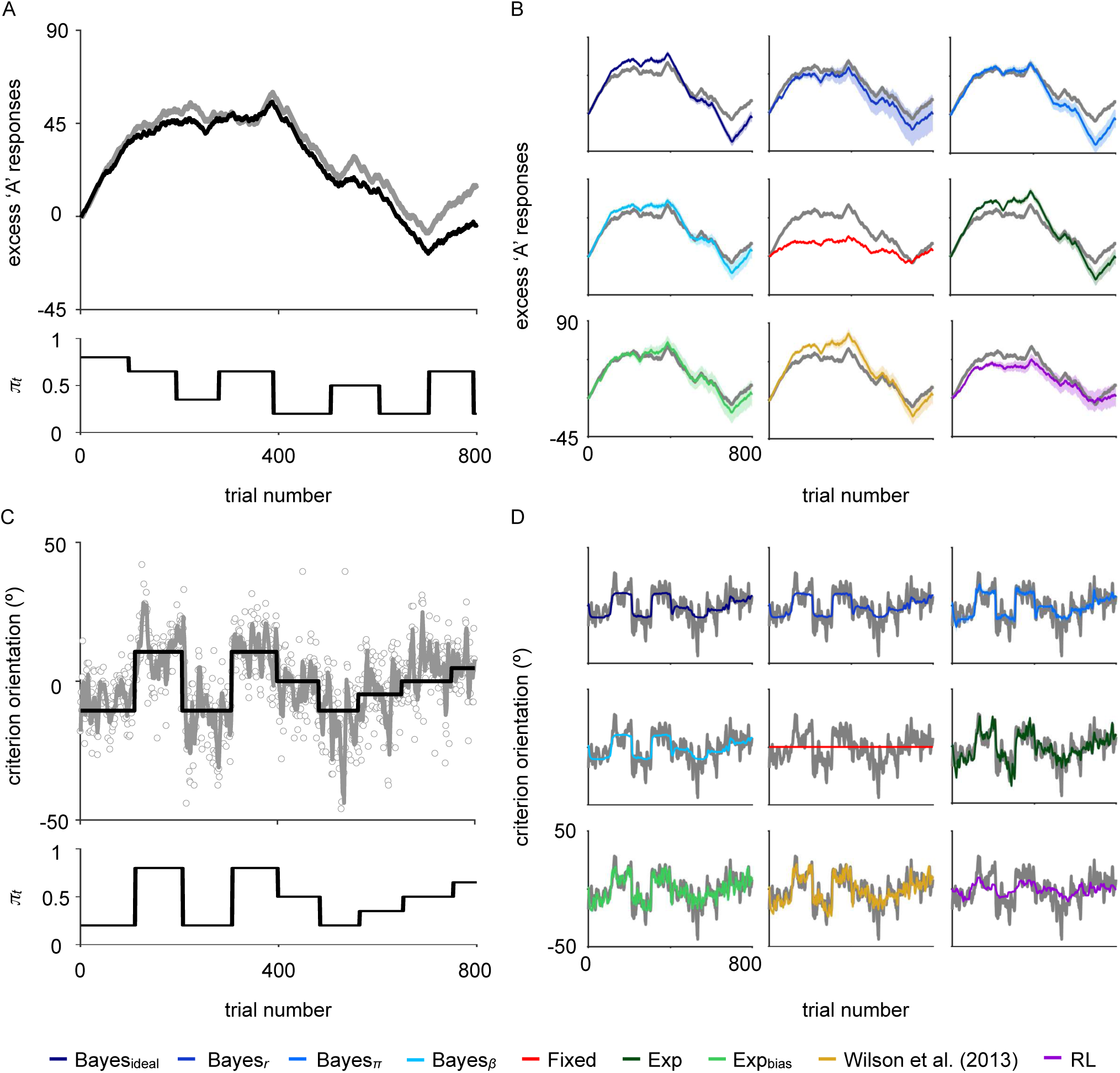
Behavioral data and model fits. A: Behavioral data from a representative observer (CWG) in the covert-criterion task. Top: The number of ‘excess’ A responses (i.e., the cumulative number of A responses minus 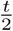) across trials. Gray line: Observer data. Black line: The true number of ‘excess’ A’s. Bottom: The corresponding category probability. B: Model fits (colored lines) for observer CWG in the covert-criterion task. Shaded regions: 68% CI for model fits. Gray: Observer data. C: Behavioral data from a representative observer (GK) in the overt-criterion task. Top: The orientation of the criterion line relative to the neutral criterion as a function of trial number. Gray circles: raw settings. Gray line: running average over a 5-trial moving window. Black line: ideal criterion for an observer with perfect knowledge of the experimental parameters. Bottom: Category probability across trials. D: Models fits (colored lines) for observer GK in the overt-criterion task. A running average was computed over a 5-trial window for visualization. Shaded regions: 68% CI on model fits. These are generally smaller than the model-fit line. Gray line: the running average computed from the observer’s data.

In the overt task, the orientation of the observer’s criterion setting, relative to the neutral criterion, is plotted as a function of trial (gray circles). For visualization, a running average was computed over a five-trial moving window (gray line). Here (Fig 2C, top), the black line represents the criterion on each trial, given perfect knowledge of the categories, sensory uncertainty, and category probability. While this is impossible for an observer to attain, we can see that the observer’s criterion follows the general trend. This suggests that observers update their criterion appropriately in response to changes in probability. That is, the criterion is set counter-clockwise from the neutral criterion when *π*_A, *t*_ > 0.5, and clockwise of neutral when *π*_A, *t*_ < 0.5. Fig 2C (bottom) shows *π*_A_ as a function of trial.

## Modeling results

### Model predictions

A qualitative comparison of the behavioral data to the ground truth suggests that observers updated their criterion in response to changes in probability. However, it does not tell us how. To explore the mechanism underlying these changes, we compared the observers’ data to multiple models. For each task and model, the mean model response for a representative subject is plotted in Fig 2B (covert task) and 2D (overt task). Shaded regions indicate 68% CIs computed from the posterior over model parameters. Specifically, we sampled parameters from the posterior over model parameters and computed the model response for the given set of parameters. We then computed the standard deviation across model responses for a large sample (see Model visualization). Shaded regions computed on overt fits are generally narrower than the data line. Qualitatively, most models captured observers’ changing criteria, with the fixed model being much worse. Differences across models are especially pronounced in the overt task. Specifically, we see that the Exp, Exp_bias_, and Wilson et al. (2013) models capture changes in criterion that occur between change points that the Bayesian models fail to capture.

### Model comparison

We quantitatively compared models by computing the log marginal likelihood (LML), also known as log Bayes factor, for each subject, model, and task. The marginal likelihood is the probability of the data given the model, marginalized over model parameters (see Methods). Here, we report differences in log marginal likelihood (ΔLML) from the best model, so that larger ΔLML correspond to worse models. We compare models both by looking at average performance across subjects (a fixed-effects analysis), and also via a Bayesian Model Selection approach (BMS; [31]) in which subjects are treated as random variables. With BMS, we estimate for each model its posterior frequency *f* in the population and its protected exceedance probability *Φ*, which is the probability that a given model is the most frequent model in the population, above and beyond chance [32].

Model comparison (Fig 3) favored the Exp_bias_ model, which outperformed the second best model Bayes_*r*_ (covert task: ΔLML = 9.27 ± 2.86; overt task: ΔLML = 8.96 ± 4.01; mean and SEM across observers) in the two tasks (covert task: *t*(10) = 3.99, *p* = 0.003; overt task: *t*(10) = 2.37, *p* = 0.04). Similarly, Bayesian model comparison performed at the group level also favored the Exp_bias_ model (covert task: *f* = 0.46 and *Φ* = 0.96; overt task: *f* = 0.38 and *Φ* = 0.84). These results suggests that observers estimate probability by taking a weighted average of recently experienced categories with a bias towards *π* = 0.5.

**Fig 3.**
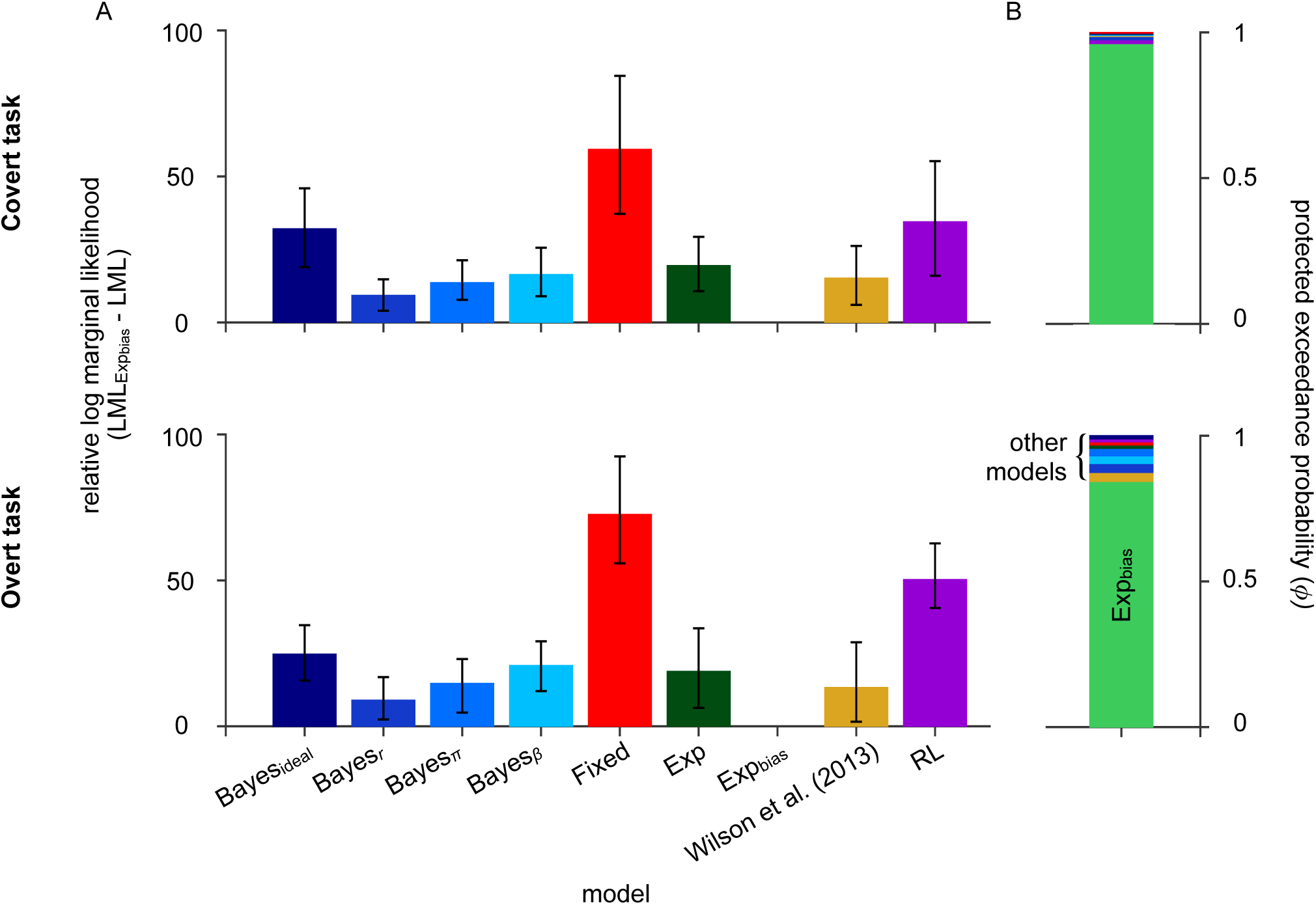
Model comparison. A: Average log marginal likelihood (LML) scores relative to the winning model, Exp_bias_ (top: covert task; bottom: overt task). Lower scores indicate a better fit. Error bars: 95% CI (bootstrapped). B: Bayesian model selection at the group level. The protected exceedance probability (*Φ*) is plotted for each model. Models are stacked in order of decreasing probability.

Fig 4A shows the number of observers that were best fit by each model for each task. To compare LML scores across tasks, and for the purpose of this analysis only, we standardized model scores for each observer and task. Standardized LML scores in the overt task are plotted as a function of standardized LML scores in the covert task in Fig 4B. We found a significant positive correlation, *r* = 0.62, *p <* 0.01, indicating that models with higher LML scores in the covert task were also higher in the overt task. This result suggests that strategy was fairly consistent across tasks. In addition, there was more variance in model scores for worse-fitting models.

**Fig 4.**
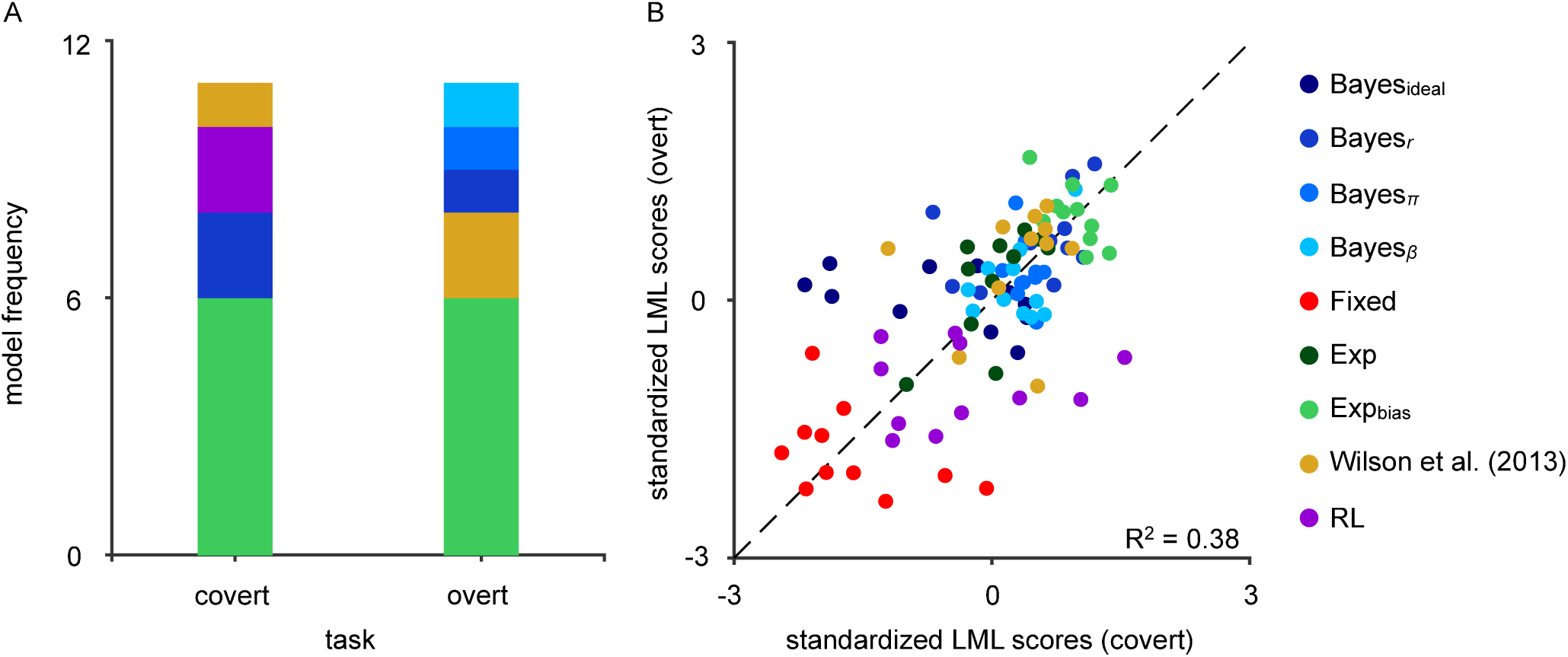
Modeling results for individual observers. A: Model frequency. The number of observers best fit by each model plotted for each task. Models are stacked in order of decreasing frequency. B: Comparison of LML scores across tasks. LML scores were standardized for each observer and task. Standardized LML scores in the overt task are plotted as a function of standardized LML scores in the covert task (colored data points). Black dashed line: identity line.

### Model parameters

We examine here the parameter estimates of the best-fitting model, Exp_bias_, recalling that the two model parameters *α*_Exp_ and *w* represent, respectively, the exponential smoothing factor and the degree to which observers exhibit conservatism. The *maximum a posteriori* (MAP) parameter estimates are plotted in Fig 5. Converting the smoothing factor to a time constant, we found that the time constant in both tasks was well below the true rate of change (covert: *τ* = [4.24, 7.18]; overt: *τ* = [3.48, 4.75]). We conducted paired-sample *t*-tests to compare the raw parameter estimates in the covert and overt tasks. We found a significant difference in *w* (*t*(10) = −2.55, *p* = 0.03), suggesting that observers were more conservative in the covert than the overt task. No significant difference was found for *α*_Exp_ (*t*(10) = −0.98, *p* = 0.35).

**Fig 5.**
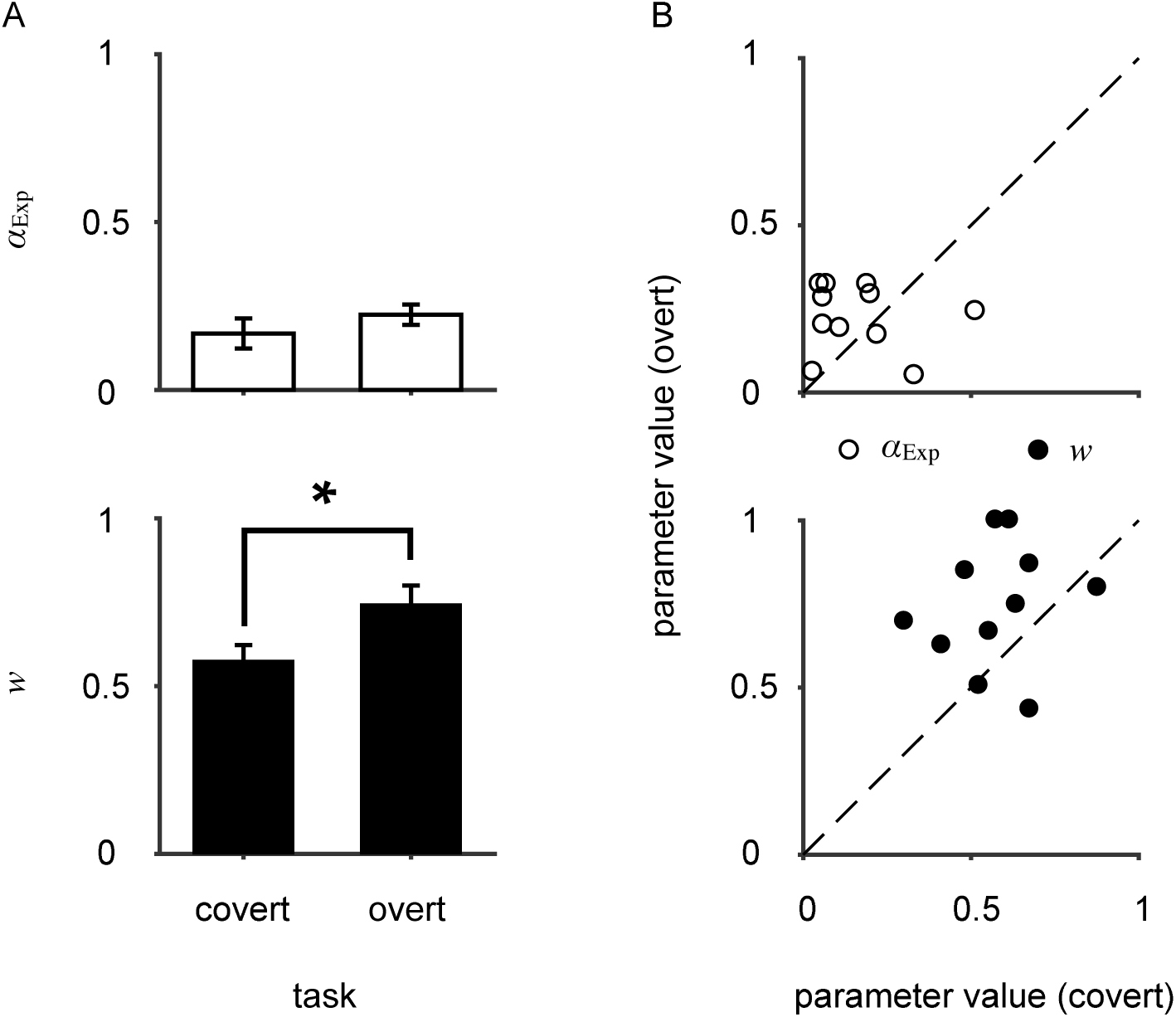
Parameter estimates for the best fitting model. A: Mean *α*_exp_ (top) and *w* (bottom) MAP values across observers. The * denotes significance at the *p <* 0.05 level. Error bars: ±SEM. B: Individual *α*_exp_ (top) and *w* (bottom) MAP values from fits in the overt task as a function of MAP values from fits in the covert task. Black dashed lines: identity line.

To investigate whether there was bias in the parameter-estimation procedure when fitting the Exp_bias_ model, we also conducted a parameter-recovery analysis. Most parameters could be recovered correctly, except for adjustment variability (*σ*_a_) in the overt task, which we found to be overestimated on average (see Section 4 in S1 Appendix for details). Note that this bias in estimating *σ*_a_ in the overt task does not affect our model comparison, which is based on LMLs and not on point estimates.

While we might expect performance to be similar across tasks and observers (i.e., a correlation between the parameter fits in each task), no significant correlations were found (*α*_Exp_: *r* = −0.14, *p* = 0.67; *w*: *r* = 0.16, *p* = 0.64). Parameter estimates for all models are shown in Tables 1 and 2.

**Table 1.** Maximum a posteriori parameter estimates ±S.E. in the covert-criterion task.

| Model | $\sigma_v$ | $r$ | $\pi$ | $\beta$ | $\alpha$ | $w$ | $l_2$ | $l_3$ | $\nu_p$ |
| --- | --- | --- | --- | --- | --- | --- | --- | --- | --- |
| Bayes<sub>ideal</sub> | $9.7 \pm 0.6$ | - | - | - | - | - | - | - | - |
| Bayes<sub>r</sub> | $9.9 \pm 0.8$ | $51.5 \pm 11.8$ | - | - | - | - | - | - | - |
| Bayes<sub>π</sub> | $10.5 \pm 0.8$ | - | $0.32 \pm 0.02$ | - | - | - | - | - | - |
| Bayes<sub>β</sub> | $10.2 \pm 0.7$ | - | - | $18.4 \pm 8.2$ | - | - | - | - | - |
| Fixed | $10.9 \pm 0.8$ | - | - | - | - | - | - | - | - |
| Exp | $9.4 \pm 0.7$ | - | - | - | $0.04 \pm 0.01$ | - | - | - | - |
| Exp<sub>bias</sub> | $10.0 \pm 0.8$ | - | - | - | $0.17 \pm 0.04$ | $0.58 \pm 0.05$ | - | - | - |
| Wilson et al. (2013) | $9.4 \pm 0.7$ | - | - | - | - | - | $12.2 \pm 2.8$ | $118.5 \pm 23.3$ | $43.3 \pm 14.4$ |
| RL | $9.2 \pm 0.7$ | - | - | - | $0.26 \pm 0.03$ | - | - | - | - |

**Table 2.**
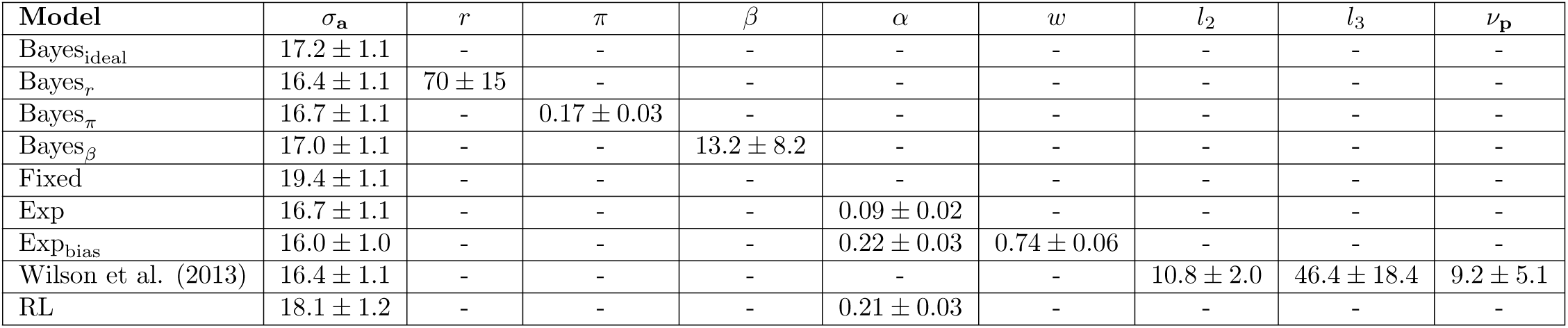
Maximum a posteriori parameter estimates ±S.E. in the overt-criterion task.

| Model | $\sigma_a$ | $r$ | $\pi$ | $\beta$ | $\alpha$ | $w$ | $l_2$ | $l_3$ | $\nu_p$ |
| --- | --- | --- | --- | --- | --- | --- | --- | --- | --- |
| Bayes<sub>ideal</sub> | $17.2 \pm 1.1$ | - | - | - | - | - | - | - | - |
| Bayes<sub>r</sub> | $16.4 \pm 1.1$ | $70 \pm 15$ | - | - | - | - | - | - | - |
| Bayes<sub>π</sub> | $16.7 \pm 1.1$ | - | $0.17 \pm 0.03$ | - | - | - | - | - | - |
| Bayes<sub>β</sub> | $17.0 \pm 1.1$ | - | - | $13.2 \pm 8.2$ | - | - | - | - | - |
| Fixed | $19.4 \pm 1.1$ | - | - | - | - | - | - | - | - |
| Exp | $16.7 \pm 1.1$ | - | - | - | $0.09 \pm 0.02$ | - | - | - | - |
| Exp<sub>bias</sub> | $16.0 \pm 1.0$ | - | - | - | $0.22 \pm 0.03$ | $0.74 \pm 0.06$ | - | - | - |
| Wilson et al. (2013) | $16.4 \pm 1.1$ | - | - | - | - | - | $10.8 \pm 2.0$ | $46.4 \pm 18.4$ | $9.2 \pm 5.1$ |
| RL | $18.1 \pm 1.2$ | - | - | - | $0.21 \pm 0.03$ | - | - | - | - |

## Discussion

Although we know that people update decision criteria in response to explicit changes in prior probability, the effects of implicit changes in prior probability on decision-making behavior are less well known. In the present study, we used model comparison to investigate the mechanisms underlying decision-making behavior in an orientation-categorization task as prior probability changed. We tested a set of models that varied in both computational and memory demands. Models were tested on data from both a covert- and overt-criterion task. A comprehensive approach, consisting of both qualitative and quantitative analysis, was performed to determine the best fitting model. We found that observers updated their decision criterion following changes in probability. Additionally, we observed systematic changes in the decision criterion during periods of stability, which was clearly evident in the overt-criterion data. Model comparison favored an exponential-averaging model with a bias towards equal probability. This suggests that observers update the decision criterion by combining on-line estimation of probability with an equal-probability prior. Ultimately, our results help explain decision-making behavior in situations in which people need to assess the probability of an outcome based on previous experience.

### Criterion updates in response to implicit changes in category probability

To determine the influence of prior probability on decision-making behavior, we examined changes in the decision criterion. First, we found that no participant was best fit by a fixed-criterion model. This finding suggests that observers update decision criteria in response to implicit changes in probability. This result is consistent with previous studies in which prior probability was explicit [2–9]. Further, this finding complements recent studies suggesting that individuals can learn and adapt to statistical regularities in changing environments [14–17, 24, 25, 33, 34]. Although this finding suggests that observers dynamically adjust decision criteria in response to changes in prior probability, it does not tell us how they do this (e.g., do observers compute on-line estimates of probability?). To uncover the mechanisms underlying changes in decision-making behavior, we compared multiple models ranging from the full Bayesian change-point detection model to a model-free reinforcement-learning (RL) model.

### Systematic criterion fluctuations

How is the decision criterion set? Qualitatively, most models appear to fit the data reasonably well in the covert task. However, when we look at data from the overt task, while Bayesian models captured the overall trend, they failed to capture local fluctuations in the decision criterion observed during periods of stability (i.e., time intervals between change points). In other words, the criterion predicted by the Bayesian models stabilized whereas the observers’ behavior did not. In contrast to the Bayesian models, the exponential models continually update the observer’s estimate of probability based on recently experienced categories. How quickly observers updated this estimate is determined in the model by the decay-rate parameter. From our model fits to the data, we found that observers had an average decay rate that was substantially smaller than the true run length distribution (4.5 vs. 100 trials, respectively), leading to frequent, systematic fluctuations in decision criteria. Although we cannot directly observe these fluctuations in the covert task, because the estimated decay rate was not significantly different across tasks we can assume the fluctuations occurred in a similar manner. Like the exponential models, the RL model was also able to capture local fluctuations in the decision criterion. However, the amplitude of the changes in criterion predicted by the RL model was generally too low compared to the data. This discrepancy was especially clear in the overt task; no participant was best fit by the RL model. These results have two important implications. (1) It is important to test alternatives to Bayesian models: observers’ behavior might be explained without requiring an internal representation of probability. (2) Using multiple tasks can provide additional insight into behavior. Here, the fluctuations in decision criteria between change points led to suboptimal behavior. Overall, our findings suggest that suboptimality arose from an incorrect, possibly heuristic inference process, that goes beyond mere sensory noise [35–37].

### A dual mechanism for criterion updating

While a number of the models captured local fluctuations in the decision criterion, we found that the Exp_bias_ model fit the data best according to a quantitative model comparison. This result suggests that observers compute on-line estimates of category probability based on recent experience. Further, the bias component of the model suggests that observers are conservative, as reflected in a long-term prior that categories are equally likely. The degree to which observers weight this prior varied across individuals and tasks. Taken together, these results suggest a dual mechanism for learning and incorporating prior probability into decisions. That is, there are (at least) two components to decision making that are acquired and updated at very different timescales.

Multiple-mechanism models have been used to describe behavior in decision-making [29] and motor behavior [38]. A model that combines delta rules predicts motor behavior better than either delta rule alone [38]. Using a combination of delta rules [29], we were able to capture the local fluctuations in criterion that the full Bayesian model missed. However, we found that a constant weight on *π* = 0.5 fit better than the multiple-node model described by Wilson and colleagues [29]. Temporal differences between their task and ours might explain some of the differences we observed, as changes occurred much more slowly in our experiment. Additionally, while fitting Wilson et al.’s model we set the hazard rate to 0.01 (the average rate of change), but observers had to learn this value throughout the experiment and may have had incorrect assumptions about the rate of change [21, 22].

### Explanations of conservatism

While we observed conservatism in both the covert- and overt-criterion tasks, we found that, on average, observers were significantly more conservative in the covert task. To understand why conservatism differs across tasks, we need to understand the differences between the tasks. While the generative model was identical across tasks, the observer’s response differed. In the covert task, observers chose between two alternatives. In the overt task, observers selected a decision criterion. This is an important difference because it allows us to potentially rule out previous explanations of conservatism, such as the use of subjective probability [4], misestimation of the relative frequency of events [39, 40], and incorrect assumptions about the sensory distributions [41, 42]; these explanations predict similar levels of conservatism across tasks. On the other hand, conservatism may be due to the use of suboptimal decision rules. Probability matching is a strategy in which participants select alternatives proportional to their probability, and has been used to explain suboptimal behavior in forced-choice tasks in which observers choose between two or more alternatives [6, 43–46]. Thus, the higher levels of conservatism in the covert task may have been due to the use of a suboptimal decision rule like probability matching, which would effectively smooth the observer’s response probability across trials. Probability matching is not applicable to responses in the overt task. Thus, the use of different decision rules may result in different levels of conservatism. These differences may also arise from an increase in uncertainty in the covert task due to less explicit feedback. An observer with greater uncertainty will rely more on the prior. Thus, conservatism may be the result of having a prior over criteria that interacts with task uncertainty. This can be tested by manipulating uncertainty over the generative model and measuring changes in conservatism. It is also possible that conservatism is the result of both the use of suboptimal decision rules and one or more of the previously proposed explanations.

### Incorrect assumptions about the generative model

While we tested a number of Bayesian models that explored an array of assumptions about the generative model, clearly one could propose even more variants. In particular, we only analyzed one such assumption at a time. A simple way to expand the model space is via a factorial comparison [36, 47], which we did not consider here due to computational intractability and the combinatorial explosion of models. Most notably, for all except the RL model we assumed knowledge of the category distributions. However, Norton et al. [16] found that for the same orientation-categorization task, category means were estimated dynamically, even after prolonged training. Similarly, Gifford et al. [48] observed suboptimality in an auditory-categorization task and found that the data were best explained by a model with non-stationary categories and prior probability that was updated using the recent history of category exemplars. This occurred despite holding categories and probability constant within a block. In fact, similar effects of non-stationarity have been observed in several other studies [49–51]. In addition to non-stationary category means, observers may also have misestimated category variance [52], especially since learning category variance takes longer than learning category means [14].

## Conclusion

In sum, our results provide a computational model for how decision-making behavior changes in response to implicit changes in prior probability. Specifically, they suggest a dual mechanism for learning and incorporating prior probability that operate at different timescales. Importantly, this helps explain behavior in situations in which assessment of probability is learned through experience. Further, our results demonstrate the need to compare multiple models and the benefit of using tasks that provide a richer, more informative dataset.

## Materials and methods

### Participants

Eleven observers participated in the experiment (mean age 26.6, range 20-31, 8 females). All observers had normal or corrected-to-normal vision. One of the observers (EHN) was also an author. The Institutional Review Board at New York University approved the experimental procedure and observers gave written informed consent prior to participation.

### Apparatus and stimuli

Stimuli were presented on a gamma-corrected Dell Trinitron P780 CRT monitor with a 31.3 × 23.8° display, a resolution of 1024 × 768 pixels, a refresh rate of 85 Hz, and a mean luminance of 40 cd/m^2^. Observers viewed the display from a distance of 54.6 cm. The experiment was programmed in MATLAB [53] using the Psychophysics Toolbox [54, 55].

Stimuli were 4.0 × 1.0° ellipses presented at the center of the display on a mid-gray background. In both the orientation-discrimination and covert-criterion tasks, trials began with a central white fixation cross (1.2°). In the overt-criterion task, a yellow line with random orientation was presented at the center of the display (5.0 × 0.5°).

### Procedure

#### Categories

In the ‘categorization’ sessions described below, stimulus orientations were drawn from one of two categories (A or B). Category distributions were Gaussian with different means (*µ*_B_ *> µ*_A_) and equal variance 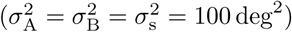. The mean of category A was chosen randomly from all possible orientations at the beginning of each session and the mean of category B was set so that *d*^′^ was approximately 1.5, which was determined for each observer based on the estimates of sensory uncertainty obtained during a ‘measurement session’ (Fig 1C).

#### Sessions

All observers participated in three 1-hour sessions. Observers completed a ‘measurement’ session first followed by two ‘categorization’ sessions. Observers completed the covert-criterion task in the first ‘categorization’ session followed by the overt-criterion task in the second, or vice versa (chosen randomly). At the beginning of each ‘categorization’ session, observers completed 200 trials of category training followed by 800 experimental trials. Prior to training, observers were provided with a detailed explanation of the category distributions and training task. After training, observers were provided with additional instructions about the subsequent ‘categorization’ task and told that the categories would remain constant for the remainder of the session but that category probability may change. Observers were not told how often category probability would change or the range of probability states.

#### Measurement task

During the ‘measurement’ session, sensory uncertainty (*σ*_v_) was estimated using a two-interval, forced-choice, orientation-discrimination task in which two black ellipses were presented sequentially on a mid-gray background. The observer reported the interval containing the ellipse that was more clockwise by keypress. Once the response was recorded, auditory feedback was provided and the next trial began. An example trial sequence is shown in Fig S3A in S1 Appendix.

The orientation of the ellipse in the first interval was chosen randomly on every trial from a uniform distribution ranging from −90 to 90°. The orientation of the second ellipse was randomly oriented clockwise or counter-clockwise of the first. The difference in orientation between the two ellipses was selected using an adaptive staircase procedure. The minimum step-size was 1° and the maximum step-size was 32°. Each observer ran two blocks. In each block, four staircases (65 trials each) were interleaved (two 1-up, 2-down and two 1-up, 3-down staircases) and randomly selected on each trial. For analyses and results see S1 Appendix.

#### Category training

Each training trial was identical to a covert-criterion trial (Fig 1A). During training there was an equal chance that a stimulus was drawn from either category. To assess learning of category distributions, observers were asked to estimate the mean orientation of each category following training. The mean of each category was estimated exactly once. The order in which category means were estimated was randomized. For estimation, a black ellipse with random orientation was displayed in the center of the display. Observers slid the mouse to the right and left to rotate the ellipse clockwise and counterclockwise, respectively and clicked the mouse to indicate they were satisfied with the setting. No feedback was provided. We computed the proportion correct for each observer to ensure category learning by comparing it to the expected proportion correct (*p*(correct) = 0.77) for *d*^′^ = 1.5. Mean estimates are plotted in Fig S4C in S1 Appendix as a function of the true category means. We computed the average estimation error for each category and observer by subtracting the estimate from the true mean. From visual inspection, it appears that training was effective with the exception of one outlier, which we assume was a lapse.

### Categorization tasks

#### Covert-criterion task

In the covert-criterion task, observers categorized ellipses based on their orientation. The start of each trial (*N*_trials_ = 800) was signaled by the appearance of a central white fixation cross (500 ms). A black oriented ellipse was then displayed at the center of the screen (300 ms). Observers categorized the ellipse as A or B by keypress. Observers received feedback as to whether they were correct on every trial. Observers received a point for every correct response and aggregate points were displayed at the top of the screen to motivate observers. In addition, the fixation cross was displayed at the center of the screen in the color corresponding to the true category (category A: green; category B: red). The next trial began immediately. An example trial sequence is depicted in Fig 1A.

#### Overt-criterion task

In the overt-criterion task, observers completed an explicit version of the categorization task described above that was developed by Norton et al. [16]. At the beginning of each trial (*N*_trials_ = 800), a line was displayed at the center of the screen. The orientation of the line was randomly selected from a uniform distribution ranging from −90 to 90°. The observers’ task was to rotate the line to indicate the criterion for that trial. Observers rotated the line clockwise or counterclockwise by sliding the mouse to the right or left and clicked the mouse to indicate their setting. Next, an ellipse was displayed under the criterion line in the color corresponding to the true category for 300 ms. Auditory feedback indicated whether the set criterion correctly categorized the ellipse. That is, observers were correct when a category A stimulus was clockwise of the criterion line or a category B stimulus was counterclockwise of the line. Observers received a point for a correct response and aggregate points were displayed at the top of the screen. The next trial began immediately. An example trial sequence is depicted in Fig 1B.

### Model fitting

For fitting, all models had one free noise parameter. In the covert-criterion task, this was sensory noise (*σ*_v_). In the overt-criterion task, sensory noise was fixed and set to the value obtained in the ‘measurement’ session, but we included a noise parameter for the adjustment of the criterion line (*σ*_a_). Fixing one noise parameter in the overt-criterion task ameliorated potential issues of lack of parameter identifiability [56], and ensured that models had the same complexity across tasks. The Bayes_ideal_ and Fixed models had no additional parameters. Each suboptimal Bayesian model had one additional parameter: Bayes_*r*_ (*r*); Bayes_*π*_ (*π*_min_); Bayes_*β*_ (*β*). The Exp and RL models also only had one additional parameter (*α*). The Exp_bias_ had two additional parameters (*α*_exp_ and *w*), and the Wilson et al. (2013) model had three (*l*_2_, *l*_3_, and *v*_p_).

To fit each model, for each subject and task we computed the logarithm of the *unnormalized* posterior probability of the parameters,

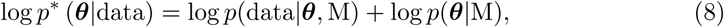

where data are category decisions in the covert-criterion task and criterion orientation in the overt-criterion task, M is a specific model, and *θ* represents the model parameters (generally, a vector). The first term of Eq 8 is the log likelihood, while the second term is the prior over parameters (see below).

We evaluated Eq 8 on a cubic grid, with bounds chosen to contain almost all posterior probability mass. The grid for all models, except Wilson et al. (2013), consisted of 100 equally spaced values for each parameter. Due to the computational demands of the Wilson et al. (2013) model, we reduced the grid to 50 equally spaced values. The grid allowed us to approximate the full posterior distribution over parameters, and also to evaluate the normalization constant for the posterior, which corresponds to the evidence or marginal likelihood, used as a metric of model comparison (see Model comparison). We reported as parameter estimates the best-fitting model parameters on the grid, that is the maximum-a-posteriori (MAP) values (see Tables 1 and 2). We used the full posterior distributions to compute posterior predictive distributions, that is, model predictions for visualization (see Model visualization), and to generate plausible parameter values for our model-recovery analysis.

#### Priors over parameters

We chose all priors to be uninformative. For noise parameters, inspired by the Jeffreys prior for scale parameters, we used uniform priors in log space over a reasonably large range ([0°, 3.4°]) [57]. For *α*_exp_, *α*_RL_, and *w* we used a uniform prior from 0 to 1. For *κ* we used a uniform prior from 2 to 200 trials. For *π*_min_ we used a uniform prior from 0 to 0.5. For *β*, we used a uniform prior on the square root of the parameter value, ranging from 0 to 10. Instead of fitting the individual nodes in the Wilson et al. (2013) model, we fit the difference between nodes, i.e., *δ*_1_ = *l*_2_ − *l*_1_ and *δ*_2_ = *l*_3_ − *l*_2_. We used a uniform prior on the square root of *δ*_1_ ranging from 1.01 to 5 and on the square root of *δ*_2_ ranging from 1.01 to 14. Finally, for *v*_p_ we used a uniform prior in log space from 0 to 5.

### Response probability

#### Covert-criterion task

For each model, parameter combination, observer, and trial in the covert-criterion task, we computed the probability of choosing category A on each trial given a stimulus, *s*_*t*_, and all previously experienced categories, *C*_1:*t* − 1_. In all models, the observer’s current decision depends on the noisy measurement, *x*_*t*_, so the probability of responding A for a given stimulus *s*_*t*_ is

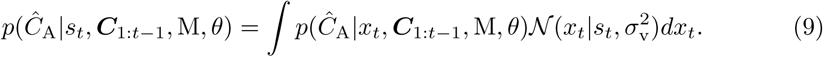

Because the current criterion setting in the RL model depends on the vector of all previous stimulus measurements, *x*_1:*t*_, the probability could not be computed analytically for this model. As an approximation, we used Monte Carlo simulations with 5000 sample measurement vectors. For each measurement vector, we applied the model’s decision rule and approximated the probability by computing the proportion of times the model chose A out of all the simulations. For all models, we included a fixed lapse rate, *λ* = 10^−4^, that is the probability of a completely random response. The probability of choosing category A, 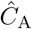, in the presence of lapses was then

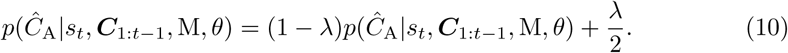

Effectively, the lapse rate acts as a regularization term that avoids excessive penalties to the likelihood of a model for outlier trials.

Next, assuming conditional independence between trials, we computed the log likelihood across all of the observer’s choices, given each model and parameter combination

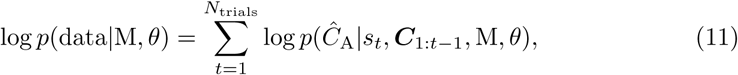

where *t* is the trial index and *N*_trials_ is the total number of trials.

#### Overt-criterion task

For each model, parameter combination, observer, and trial in the overt-criterion task, we computed the decision criterion on each trial. For all models except the RL model, the criterion was computed as in S1 Appendix. For the RL model, the criterion was computed as in Eq 7 for 5000 sample measurement vectors. For all models in the overt-criterion task, the criterion was corrupted by adjustment noise with variance 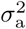, so that 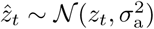, where *z*_*t*_ was the observer’s chosen criterion at trial *t*, and 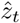 was the actual reported criterion after adjustment noise. In addition, the observer had a chance of lapsing (e.g., a misclick), in which case the response was uniformly distributed in the range. Therefore, the probability that the observer reports the criterion 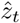 was

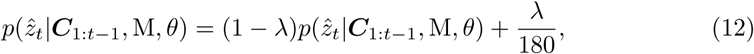

with *λ* = 5 × 10^−5^. As in the covert-criterion task, we computed the log likelihood across all trials by summing the log probability

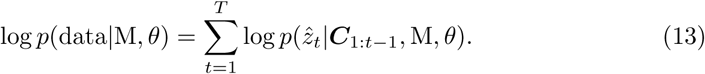

### Model comparison

To obtain a quantitative measure of model fit, for each observer, model, and task we computed the log marginal likelihood (LML) by integrating over the parameters in

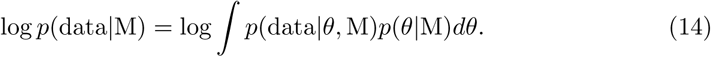

To approximate the integral in Eq 14, we marginalized across each parameter dimension using the trapezoidal method. Assuming equal probability across models, the marginal likelihood was proportional to the posterior probability of a given model and thus represents a principled metric for comparison that automatically accounts for both goodness of fit and model complexity via Bayesian Occam’s razor [58]. Penalizing for model complexity is a desirable feature of a model-comparison metric to reduce overfitting.

In addition to the Bayesian model-comparison metric described above, we computed the Akaike Information Criterion (AIC) [59] for each of our models. AIC is one of many information criteria that penalize the maximum log likelihood by a term that increases with the number of parameters. LML and AIC results were consistent (see S1 Appendix for model comparison using AIC scores).

For comparison purposes, we report relative model-comparison scores, ΔLML and ΔAIC. We used bootstrapping to compute confidence intervals on the mean difference scores. Specifically, we simulated 10,000 sample data sets. For each simulated dataset we sampled, with replacement, 11 difference scores (the same number of difference scores as observers) and calculated the mean. To determine the 95% CI, we sorted the mean difference scores and determined the scores that corresponded to the 2.5 and 97.5 percentiles.

For an additional analysis at the group level, we used the random-effects Bayesian model selection analysis (BMS) developed by Stephan et al. [31] and expanded on by Rigoux et al. [32]. Specifically, using observers’ LML scores we computed the protected exceedance probability *Φ* and the posterior model frequency for each model. Exceedance probability represents the probability that one model is the most frequent decision-making strategy in the population, given the group data, above and beyond chance. This analysis was conducted using the open-source software package Statistical Parametric Mapping (SPM12; http://www.fil.ion.ucl.ac.uk/spm).

### Model visualization

For each model, observer, and task, we randomly sampled 1000 parameter combinations from the joint posterior distribution with replacement. For each parameter combination, we simulated model responses using the same stimuli that were presented to the observer. Because the model output in the covert task was the probability of reporting category A, for each trial in a simulated dataset we simulated 10,000 model responses (i.e., category decisions), calculated the cumulative number of A’s for each simulated dataset, and averaged the results. The mean and standard deviation were computed across all simulated datasets in both tasks. Model fit plots show the mean response (colored line) with shaded regions representing one standard deviation from the mean. Thus, shaded regions represent a 68% confidence interval on model fits.

### Model recovery

To ensure that our models were discriminable, we performed a model-recovery analysis, details of which can be found in S1 Appendix. In addition to the model-recovery analysis, we also performed a parameter-recovery analysis for the winning model (Exp_bias_). This was done to determine whether our parameter estimation procedure was biased for each parameter and task.

## Supporting information

### S1 Appendix. Supplementary Information

Ideal observer model derivation; additional models; model comparison with AIC; model recovery analysis; measurement task; category training.

## Supporting information

## Acknowledgments

We would like to thank Chris Grimmick for helping with data collection. This work utilized the NYU IT High Performance Computing resources and services.

