## Supplementary material for "Human online adaptation to changes in prior probability"

November 28, 2018

### 1 Ideal observer model derivation

Our derivation of the Bayesian online change-point detection algorithm for the ideal observer generalizes that of Adams and MacKay [1]. For clarity and ease of reference, we report here the full derivation; only the broad outline is described in the main text.

#### 1.1 Bayesian online change-point detection

The Bayesian observer estimates the posterior distribution over the current *run length*, or time since the last change point, and the state (category probabilities) before the last change point, given the data (category labels) observed so far. We denote the length of the run at the end of trial $t$ by $r_t$. Similarly, we denote with $\pi_t$ and $\xi_t$ the current state and the state before the last change point, both measured at the end of trial $t$. Here, $\pi_t$ represents the probability that, on the subsequent trial, the category will be A (the probability of category B is $1 - \pi_t$). Both $\pi_t, \xi_t \in S_\pi$, where $S_\pi$ is a discrete set of possible states. In the experiment, $S_\pi = \{0.2, 0.35, 0.5, 0.65, 0.8\}$. We use the notation $\boldsymbol{C}_t^{(r)}$ to indicate the set of observations (category labels) associated with the run $r_t$, which is $\boldsymbol{C}_{t-r_t+1:t}$ for $r_t > 0$, and $\emptyset$ for $r_t = 0$. We use the subscript colon operator $\boldsymbol{C}_{t':t}$ to denote the sub-vector of $\boldsymbol{C}$ (the full sequence of observed categories) with elements from $t'$ to $t$ included.

We write the predictive distribution of category by marginalizing over run lengths $r_t$ and previous states $\xi_t$,

$$P(C_{t+1}|\boldsymbol{C}_{1:t}) = \sum_{r_t} \sum_{\xi_t} P(C_{t+1}|r_t, \xi_t, \boldsymbol{C}_t^{(r)}) P(r_t, \xi_t|\boldsymbol{C}_{1:t}). \tag{S1}$$

We assume that, in the case of a change point at the end of trial $t$, the new state might have Markovian dependence on the previous state, that is

$\pi_t \sim P(\pi_t|\pi_{t-1})$. This is a generalization of the model of Adams and MacKay [1], in which the distribution parameters were assumed to be independent after change points. In the experiment, $P(\pi_t|\pi_{t-1}) = \frac{1}{|S_\pi|-1}[\![\pi_t \neq \pi_{t-1}]\!]$. We use $[\![A]\!]$ to denote *Iverson's bracket* which is 1 if the expression $A$ is true, and 0 otherwise [2].

To find the posterior distribution (the second term in Eq S1)

$$P(r_t, \xi_t|\boldsymbol{C}_{1:t}) = \frac{P(r_t, \xi_t, \boldsymbol{C}_{1:t})}{P(\boldsymbol{C}_{1:t})} \tag{S2}$$

we write the joint distribution over run length, previous state and observed data recursively,

$$\begin{aligned} P(r_t, \xi_t, \boldsymbol{C}_{1:t}) &= \sum_{r_{t-1}} \sum_{\xi_{t-1}} P(r_t, r_{t-1}, \xi_t, \xi_{t-1}, \boldsymbol{C}_{1:t}) \\ &= \sum_{r_{t-1}} \sum_{\xi_{t-1}} P(r_t, \xi_t, C_t|r_{t-1}, \xi_{t-1}, \boldsymbol{C}_{1:t-1}) P(r_{t-1}, \xi_{t-1}, \boldsymbol{C}_{1:t-1}) \\ &= \sum_{r_{t-1}} \sum_{\xi_{t-1}} P(r_t, \xi_t|r_{t-1}, \xi_{t-1}, \boldsymbol{C}_t^{(r)}) \\ &\qquad \times P(C_t|r_{t-1}, \xi_{t-1}, \boldsymbol{C}_{t-1}^{(r)}) P(r_{t-1}, \xi_{t-1}, \boldsymbol{C}_{1:t-1}). \end{aligned} \tag{S3}$$

Note that the justification for specializing from $\boldsymbol{C}_{1:t}$ to $\boldsymbol{C}_t^{(r)}$ and $\boldsymbol{C}_{t-1}^{(r)}$ will become clear in the derivations below. We can rewrite Eq S3 in terms of the posterior distribution as

$$\begin{aligned} P(r_t, \xi_t|\boldsymbol{C}_{1:t}) &= \frac{1}{P(\boldsymbol{C}_{1:t})} P(r_t, \xi_t, \boldsymbol{C}_{1:t}) \\ &= \frac{P(\boldsymbol{C}_{1:t-1})}{P(\boldsymbol{C}_{1:t})} \sum_{r_{t-1}} \sum_{\xi_{t-1}} P(r_t, \xi_t|r_{t-1}, \xi_{t-1}, \boldsymbol{C}_t^{(r)}) \\ &\qquad \times P(C_t|r_{t-1}, \xi_{t-1}, \boldsymbol{C}_{t-1}^{(r)}) P(r_{t-1}, \xi_{t-1}|\boldsymbol{C}_{1:t-1}). \end{aligned} \tag{S4}$$

Eq S4 is the basis for the iterative Bayesian algorithm, since it allows us to derive the posterior distribution at time $t$ as a function of the posterior distribution and a number of auxiliary variables at time $t-1$.

For computational convenience, we rewrite the posterior from Eq S4 as an unnormalized posterior

$$U(r_t, \xi_t|\boldsymbol{C}_{1:t}) = \sum_{r_{t-1}} \sum_{\xi_{t-1}} P(r_t, \xi_t|r_{t-1}, \xi_{t-1}, \boldsymbol{C}_t^{(r)}) P_t^{(r_{t-1}, \xi_{t-1})} \tag{S5}$$

where we introduced $P_t^{(r_{t-1},\xi_{t-1})}$ to denote the posterior from the previous trial times the conditional predictive probability for the current category,

$$P_t^{(r_{t-1},\xi_{t-1})} \equiv P(C_t|r_{t-1},\xi_{t-1},\boldsymbol{C}_{t-1}^{(r)})P(r_{t-1},\xi_{t-1}|\boldsymbol{C}_{1:t-1}). \tag{S6}$$

To compute the unnormalized posterior in Eq S5, we need:

- the conditional predictive probability, which we compute in the following Section 1.1.1;
- the change-point posterior, which we compute in Section 1.1.2;
- the posterior from the previous trial.

We put everything together in Section 1.1.3.

##### 1.1.1 Conditional predictive probability

The posterior over state at the end of trial $t-1$, given the last $r_{t-1}$ trials and the previous state $\xi_{t-1}$, is

$$P(\pi_{t-1}|r_{t-1},\xi_{t-1},\boldsymbol{C}_{t-1}^{(r)}) \propto [\![\pi_{t-1} \neq \xi_{t-1}]\!] P(\pi_{t-1}|r_{t-1},\boldsymbol{C}_{t-1}^{(r)}). \tag{S7}$$

For computational convenience, we denote $\Psi_t^{(r,\pi)} \equiv P(\pi_t = \pi|r_t = r, \boldsymbol{C}_t^{(r)})$ and we store it in a table. Clearly, $\Psi_t^{(r,\pi)}$ depends only on the length of the run $r$, the category probability $\pi$ and the number of times category A occurs during the run. In the algorithm below, $\Psi$ is computed iteratively trial-by-trial and values of $\Psi$ are computed only for combinations of run length and number of A categories that occur in the sequence. The conditional predictive probability for observing $C_t$, using Eq S7, is

$$\begin{aligned} P(C_t|r_{t-1},\xi_{t-1},\boldsymbol{C}_{t-1}^{(r)}) &= \sum_{\pi_{t-1}} P(C_t|\pi_{t-1})P(\pi_{t-1}|r_{t-1},\xi_{t-1},\boldsymbol{C}_{t-1}^{(r)}) \\ &\propto \sum_{\pi_{t-1}} P(C_t|\pi_{t-1})[\![\pi_{t-1} \neq \xi_{t-1}]\!]\Psi_{t-1}^{(r_{t-1},\pi_{t-1})}. \end{aligned} \tag{S8}$$

##### 1.1.2 The change-point posterior

The conditional posterior on the change point (that is, run length) and previous state, $P(r_t, \xi_t|r_{t-1},\xi_{t-1},\boldsymbol{C}_t^{(r)})$, has a sparse representation since it has only two outcomes: the run length either continues to grow ($r_t = r_{t-1} + 1$ and $\xi_t = \xi_{t-1}$) or a change point occurs ($r_t = 0$ and the posterior over $\xi_t$ is the posterior over $\pi_{t-1}$ computed from $\boldsymbol{C}_t^{(r)}$). We have

$$P(r_t, \xi_t | r_{t-1}, \xi_{t-1}, \boldsymbol{C}_t^{(r)}) = P(r_t | r_{t-1}) P(\xi_t | r_t, r_{t-1}, \xi_{t-1}, \boldsymbol{C}_t^{(r)}). \tag{S9}$$

The probability of a run length after a change point is

$$P(r_t | r_{t-1}) = \begin{cases} H(r_{t-1} + 1) & \text{if } r_t = 0 \\ 1 - H(r_{t-1} + 1) & \text{if } r_t = r_{t-1} + 1 \\ 0 & \text{otherwise} \end{cases} \tag{S10}$$

where the function $H(\tau)$ is the *hazard function*,

$$H(\tau) = \frac{P_{\text{gap}}(g = \tau)}{\sum_{t=\tau}^{\infty} P_{\text{gap}}(g = t)}, \text{ for } \tau \geq 1. \tag{S11}$$

$P_{\text{gap}}$ is the prior over run lengths. In our experimental setup, $P_{\text{gap}}(g) = \frac{1}{g_{\max} - g_{\min} + 1} [\![ g_{\min} \leq g \leq g_{\max} ]\!]$, with $g_{\min} = 80$ and $g_{\max} = 120$.

The conditional posterior over the previous state is

$$P(\xi_t | r_t, r_{t-1}, \xi_{t-1}, C_t, \boldsymbol{C}_{t-1}^{(r)}) = \begin{cases} P(\pi_{t-1} = \xi_t | r_{t-1}, \xi_{t-1}, C_t, \boldsymbol{C}_{t-1}^{(r)}) & \text{if } r_t = 0 \\ \delta(\xi_t - \xi_{t-1}) & \text{otherwise} \end{cases} \tag{S12}$$

where $P(\pi_{t-1} | r_{t-1}, \xi_{t-1}, C_t, \boldsymbol{C}_{t-1}^{(r)})$ is the posterior over state given that $C_t$ has just been observed,

$$\begin{aligned} P(\pi_{t-1} | r_{t-1}, \xi_{t-1}, C_t, \boldsymbol{C}_{t-1}^{(r)}) &\propto P(\pi_{t-1}, r_t, r_{t-1}, \xi_{t-1}, C_t, \boldsymbol{C}_{t-1}^{(r)}) \\ &\propto P(C_t | \pi_{t-1}) P(\pi_{t-1} | r_{t-1}, \xi_{t-1}, \boldsymbol{C}_{t-1}^{(r)}) P(r_t | r_{t-1}) \\ &\propto P(C_t | \pi_{t-1}) [\![ \pi_{t-1} \neq \xi_{t-1} ]\!] \Psi_{t-1}^{(r_{t-1}, \pi_{t-1})} \end{aligned} \tag{S13}$$

where in the last step $P(r_t | r_{t-1})$ is constant in $\pi_{t-1}$ and therefore irrelevant.

##### 1.1.3 Iterative posterior update and boundary conditions

Using Eqs S5 and S9, the iterative posterior update equation becomes

$$U(r_t, \xi_t | \boldsymbol{C}_{1:t}) = \sum_{r_{t-1}} P(r_t | r_{t-1}) \sum_{\xi_{t-1}} P(\xi_t | r_{t-1}, \xi_{t-1}, \boldsymbol{C}_t^{(r)}) P_t^{(r_{t-1}, \xi_{t-1})} \tag{S14}$$

which is computed separately for the case $r_t = 0$ and $r_t > 0$ via Eqs S7-S13.

We assume as boundary conditions that a change point just occurred and uniform probability across previous states

$$P(r_0, \xi_0|\emptyset) = \delta(r_0)\frac{1}{|S_\pi|}. \tag{S15}$$

Once we have $U(r_t, \xi_t|\boldsymbol{C}_{1:t})$, we can easily obtain the *normalized* posterior $P(r_t, \xi_t|\boldsymbol{C}_{1:t})$ by computing the normalization constant via a discrete summation over run lengths $r_t$ and previous states $\xi_t$,

$$P(r_t, \xi_t|\boldsymbol{C}_{1:t}) = \frac{U(r_t, \xi_t|\boldsymbol{C}_{1:t})}{\mathcal{Z}(\boldsymbol{C}_{1:t})}, \quad \text{with } \mathcal{Z}(\boldsymbol{C}_{1:t}) = \sum_{r_t}\sum_{\xi_t} U(r_t, \xi_t|\boldsymbol{C}_{1:t}). \tag{S16}$$

This result together with Eq S1 allows the observer to compute $P(C_{t+1}|\boldsymbol{C}_{1:t})$.

#### 1.2 Task-dependent predictive distributions

Armed with an expression for the observer's posterior distribution over run lengths and previous states, given all trials experienced so far (Eq S16), we can now compute the predictive distributions for the observer's response at trial $t$ for the covert- and overt-criterion tasks.

##### 1.2.1 Covert-criterion task

The probability density of a noisy measurement $x_t$ is

$$p(x_t|C_t) = \mathcal{N}(x_t|\mu_{C_t}, \sigma^2) \tag{S17}$$

with $\sigma^2 \equiv \sigma_\mathrm{v}^2 + \sigma_s^2$, where $\sigma_\mathrm{v}^2$ is the observer's visual measurement noise and $\sigma_s^2$ is the stimulus variance. The conditional posterior for category $C_t$, after observing $x_t$, is

$$P(C_t|x_t, \boldsymbol{C}_{1:t-1}) = \frac{P(x_t|C_t)P(C_t|\boldsymbol{C}_{1:t-1})}{\sum_{C'_t} P(x_t|C'_t)P(C'_t|\boldsymbol{C}_{1:t-1})}. \tag{S18}$$

We assume that for a given noisy measurement the observer responds $\hat{C}_t$ if that category is more probable, that is,

$$P(\hat{C}_t|x_t, \boldsymbol{C}_{1:t-1}) = [\![P(C_t|x_t, \boldsymbol{C}_{1:t-1}) > 0.5]\!]. \tag{S19}$$

The probability of observing response $\hat{C}_t$ for a given stimulus $s_t$ is therefore

$$P(\hat{C}_t|s_t, \boldsymbol{C}_{1:t-1}) = \int P(\hat{C}_t|x_t, \boldsymbol{C}_{1:t-1}) \mathcal{N}(x_t|s_t, \sigma_{\text{v}}^2) dx_t \tag{S20}$$

which can be easily computed via 1-D numerical integration over a grid of regularly spaced $x_t$ using trapezoidal or Simpson's rule [3].

We can also consider an observer model with lapses that occasionally reports the wrong category with probability $0 \leq \lambda \leq 1$,

$$P_{\text{lapse}}(\hat{C}_t|s_t, \boldsymbol{C}_{1:t-1}) = (1-\lambda) P(\hat{C}_t|s_t, \boldsymbol{C}_{1:t-1}) + \frac{\lambda}{|C|}, \tag{S21}$$

where $|C|$ is the number of categories in the task ($|C| = 2$ in our case), and we assume equal response probability across categories for lapses.

##### 1.2.2 Overt-criterion task

The optimal criterion $z_{opt}$ is the point at which $P(C_{\text{A}}|x, t) = P(C_{\text{B}}|x, t)$, given the available information at trial $t$. Specifically, noting that $P(C, \pi_t|x) \propto P(x|C)P(C|\pi_t)P(\pi_t)$, we have

$$\begin{aligned}
\sum_{\pi_t} P(x|C_{\text{A}})P(C_{\text{A}}|\pi_t)P(\pi_t) &= \sum_{\pi_t} P(x|C_{\text{B}})P(C_{\text{B}}|\pi_t)P(\pi_t) \\
P(x|C_{\text{A}}) \sum_{\pi_t} \pi_t P(\pi_t) &= P(x|C_{\text{B}}) \sum_{\pi_t} (1-\pi_t) P(\pi_t) \\
\frac{\sum_{\pi_t} \pi_t P(\pi_t)}{\sum_{\pi_t} (1-\pi_t) P(\pi_t)} &= e^{-\frac{(x-\mu_{\text{B}})^2}{2\sigma^2} + \frac{(x-\mu_{\text{A}})^2}{2\sigma^2}} \\
\Longrightarrow z_t^{\text{opt}} &= \frac{\sigma^2 \log \Gamma_t}{\mu_{\text{B}} - \mu_{\text{A}}} + \frac{1}{2}(\mu_{\text{A}} + \mu_{\text{B}})
\end{aligned} \tag{S22}$$

where we have defined $\Gamma_t = \frac{\sum_{\pi_t} \pi_t P(\pi_t)}{\sum_{\pi_t} (1-\pi_t) P(\pi_t)}$. We assume that $\mu_{\text{A}}$ and $\mu_{\text{B}}$ are known exactly from the training session.

The probability that the observer reports criterion $\hat{z}_t$ at trial $t$ is

$$P(\hat{z}_t|\boldsymbol{C}_{1:t-1}) = \mathcal{N}(\hat{z}_t|z_t^{\text{opt}}, \sigma_{\text{a}}^2), \tag{S23}$$

where $\sigma_{\text{a}}^2$ is criterion-placement (adjustment) noise. Note that the likelihood at trial $t$ is based on information gathered through trial $t-1$.

We can also consider an observer model with lapses who occasionally reports a criterion uniformly at random with probability $0 \leq \lambda \leq 1$,

$$P_{\text{lapse}}(\hat{z}_t|\boldsymbol{C}_{1:t-1}) = (1-\lambda)P(\hat{z}_t|\boldsymbol{C}_{1:t-1}) + \frac{\lambda}{180}. \tag{S24}$$

#### 1.3 Algorithm

In the following, we use the notation $P(\boldsymbol{x}|\boldsymbol{y}) \propto f(\boldsymbol{x},\boldsymbol{y})$ to indicate that the user needs to compute $f(\boldsymbol{x},\boldsymbol{y})$ and then normalize as follows, $P(\boldsymbol{x}|\boldsymbol{y}) = \frac{f(\boldsymbol{x},\boldsymbol{y})}{\sum_{\boldsymbol{x}'} f(\boldsymbol{x}',\boldsymbol{y})}$.

1. Initialize
   (a) Posterior $P(r_0, \xi_0|\emptyset) = \delta(r_0)\frac{1}{|S_\pi|}$
   (b) Lookup table $\Psi^{(r_0,\pi_0)} = \delta(r_0)\frac{1}{|S_\pi|}$
   (c) Set trial $t = 1$
2. Observe new category $C_t$
3. Compute auxiliary variables
   (a) Evaluate predictive probability (Eq S8)
   $$P(C_t|r_{t-1}, \xi_{t-1}, \boldsymbol{C}^{(r)}_{t-1}) \propto \sum_{\pi_{t-1}} P(C_t|\pi_{t-1})[\![\pi_{t-1} \neq \xi_{t-1}]\!]\Psi^{(r_{t-1},\pi_{t-1})}$$
   (b) Evaluate the predictive probability times posterior probability (Eq S6)
   $$P_t^{(r_{t-1},\xi_{t-1})} = P(C_t|r_{t-1}, \xi_{t-1}, \boldsymbol{C}^{(r)}_{t-1})P(r_{t-1}, \xi_{t-1}|\boldsymbol{C}_{1:t-1})$$
   (c) Evaluate the posterior probability over state (from Eq S13)
   $$P(\pi_{t-1}|r_{t-1}, \xi_{t-1}, C_t, \boldsymbol{C}^{(r)}_{t-1}) \propto P(C_t|\pi_{t-1})[\![\pi_{t-1} \neq \xi_{t-1}]\!]\Psi^{(r_{t-1},\pi_{t-1})}$$
4. Update run length and previous-state posterior

(a) Calculate the unnormalized change-point probabilities (Eq S14)

$$U(r_t = 0, \xi_t | \boldsymbol{C}_{1:t}) = \sum_{r_{t-1}} H(r_{t-1}+1) \times \sum_{\xi_{t-1}} P(\pi_{t-1} = \xi_t | r_{t-1}, \xi_{t-1}, C_t, \boldsymbol{C}_{t-1}^{(r)}) P_t^{(r_{t-1}, \xi_{t-1})}$$

(b) Calculate the unnormalized growth probabilities (see Eq S14)

$$U(r_t = r_{t-1}+1, \xi_t = \xi_{t-1} | \boldsymbol{C}_{1:t}) = [1 - H(r_{t-1}+1)]\, P_t^{(r_{t-1}, \xi_{t-1})}$$

(c) Calculate the normalization (Eq S16)

$$\mathcal{Z}(\boldsymbol{C}_{1:t}) = \sum_{r_t} \sum_{\xi_t} U(r_t, \xi_t | \boldsymbol{C}_{1:t})$$

(d) Determine the posterior distribution (Eq S16)

$$P(r_t, \xi_t | \boldsymbol{C}_{1:t}) = \frac{U(r_t, \xi_t | \boldsymbol{C}_{1:t})}{\mathcal{Z}(\boldsymbol{C}_{1:t})}$$

5. Bookkeeping and predictions

(a) Update sufficient statistics for all $r$ and $\pi$

$$\tilde{\Psi}_t^{(r,\pi)} = \begin{cases} 1 & \text{if } r = 0 \\ \Psi_{t-1}^{(r,\pi)} P(C_t | \pi_{t-1} = \pi) & \text{if } r > 0 \end{cases}$$

$$\Psi_t^{(r,\pi)} = \frac{\tilde{\Psi}_t^{(r,\pi)}}{\sum_{\pi\prime} \tilde{\Psi}_t^{(r,\pi\prime)}}$$

(b) Compute the predictive distribution of category (Eq S1)

(c) Store predictive posterior over $\pi_t$

$$P(\pi_t | \boldsymbol{C}_{1:t}) \propto \sum_{r_t} \sum_{\xi_t} \Psi_t^{(r_t, \pi_t)} [\![\pi_t \neq \xi_t]\!] P(r_t, \xi_t | \boldsymbol{C}_{1:t})$$

6. Increase trial index $t \leftarrow t+1$ and return to step 2

For each trial $t$, the posterior predictive distributions calculated in steps 5b and 5c are used to compute the observer's response probabilities in the covert- and overt-criterion tasks, respectively, as described in Section 1.2.

### 2 Additional models

We describe here a number of model variants which we did not include in the main text for reasons of space. For all additional models, we report as a model comparison metric the difference in log marginal likelihood ($\Delta$LML) with respect to a baseline model (mean $\pm$ SEM across subjects). Usually, unless stated otherwise, we take as baseline the best-fitting model described in the main text, $\text{Exp}_{\text{bias}}$. Positive values of $\Delta$LML denote a worse-fitting model than baseline.

#### 2.1 Bayesian

The main text discusses four Bayesian models. $\text{Bayes}_{\text{ideal}}$ is the algorithm above, using the precise generative model for our experiment. $\text{Bayes}_{r}$ uses the same algorithm, but adds a free parameter for the run-length distribution (and hence the hazard function) assumed by the observer. $\text{Bayes}_{\pi}$ also uses the same algorithm, but adds a parameter for the range of the set of five states assumed by the observer. $\text{Bayes}_{\beta}$ assumes the observer uses a beta-distributed prior over states. To implement this observer requires minor modifications of the algorithm. In particular, the beta prior is substituted for the uniform distribution in the initialization steps for $P(r_0, \xi_0|\emptyset)$ and $\Psi^{(r_0,\pi_0)}$ as well as the update step for $\tilde{\Psi}_t^{(r,\pi)}$ in the case where $r_t = 0$.

To ensure the robustness of our results we fit two additional suboptimal Bayesian models and compared each model to the winning model ($\text{Exp}_{\text{bias}}$). To capture conservatism as we did in the $\text{Exp}_{\text{bias}}$ model, we fit a model that took a weighted average between the probability predicted by the ideal observer model and $\pi = 0.5$ ($\text{Bayes}_{\text{bias}}$). The weight on the probability computed by an ideal observer was defined by the parameter $w$ with range $0 \leq w \leq 1$, such that 0 indicated the use of a fixed criterion and 1 the optimal. This model is similar to the $\text{Bayes}_{\beta}$ model described in the main text with a symmetric hyperprior on $\pi$, in that both result in conservatism. However, we ran it to ensure that the fits did not change when the parameterization was identical to the $\text{Exp}_{\text{bias}}$ model. We also fit a three-parameter model, in which the maximum run length $r$ and the hyperparameter $\beta$ were both free parameters ($\text{Bayes}_{r,\beta}$). We chose these parameters because the $\text{Bayes}_{r}$ model was the best fitting Bayesian model tested, and the $\text{Bayes}_{\beta}$ model takes into account conservatism, which we observed in our data. Neither of these additional models fit better than the $\text{Exp}_{\text{bias}}$ ($\text{Bayes}_{\text{bias}}$: $\Delta\text{LML}_{\text{covert}} =$

$19.92 \pm 5.00$, $\Delta\text{LML}_{\text{overt}} = 20.49 \pm 3.43$; $\text{Bayes}_{r,\beta}$: $\Delta\text{LML}_{\text{covert}} = 5.89 \pm 2.10$, $\Delta\text{LML}_{\text{overt}} = 10.54 \pm 4.32$).

#### 2.2 Reinforcement learning – probability updating

The following model ($\text{RL}_{\text{prob}}$) differs from the RL models in the main text in that it updates the category probability (as opposed to updating the decision criterion), which makes it similar to the exponential-averaging model. Similarly to the RL models in the main text (and in contrast with the Exp models), it updates probability according to a delta rule which is applied only after *incorrect* responses. After each response at trial $t$, the probability estimate for the next trial is updated using the following delta rule,

$$\hat{\pi}_{\text{A},t+1} = \begin{cases} \hat{\pi}_{\text{A},t} & \text{if correct} \\ \hat{\pi}_{\text{A},t} + \alpha_{\text{prob}}(C_t - \hat{\pi}_{\text{A},t}) & \text{if incorrect} \end{cases} \tag{S25}$$

where $\hat{\pi}_{\text{A},t}$ is the observer's estimate of the probability for category A on trial $t$ ($\hat{\pi}_{\text{B},t} = 1 - \hat{\pi}_{\text{A},t}$), $\alpha_{\text{prob}}$ is the learning rate, and $C_t$ is the current category label. Thus, the probability estimate is updated when negative feedback is received by taking a small step in the direction of the most recently experienced category. This model has two free parameters ($\alpha_{\text{prob}}$ and either $\sigma_{\text{v}}$ or $\sigma_{\text{a}}$).

To capture conservatism, we considered an additional model ($\text{RL}_{\text{prob, bias}}$) in which we took a weighted average between the probability estimate and $\pi = 0.5$, which added another free parameter $w$ to the model.

In terms of model comparison, both models were indistinguishable from the fixed criterion model ($\text{RL}_{\text{prob}}$: $\Delta\text{LML}_{\text{covert}} = -2.48 \pm 2.11$, $\Delta\text{LML}_{\text{overt}} = -2.92 \pm 1.22$; $\text{RL}_{\text{prob, bias}}$: $\Delta\text{LML}_{\text{covert}} = 0.36 \pm 2.18$, $\Delta\text{LML}_{\text{overt}} = 1.09 \pm 1.34$). Furthermore, the fits were significantly worse than the $\text{Exp}_{\text{bias}}$ model ($\text{RL}_{\text{prob}}$: $\Delta\text{LML}_{\text{covert}} = 61.75 \pm 13.44$, $\Delta\text{LML}_{\text{overt}} = 75.53 \pm 10.20$; $\text{RL}_{\text{prob, bias}}$: $\Delta\text{LML}_{\text{covert}} = 58.91 \pm 13.71$, $\Delta\text{LML}_{\text{overt}} = 71.52 \pm 9.91$).

#### 2.3 Wilson et al. (2013)

The Wilson et al. (2013) model was developed as an approximation to the full change-point detection model. Their approximation used a mixture of delta rules, each of which alone is identical to our Exp model with different learning rates. In the main text, we fit a three node model with two free node parameters ($l_2$ and $l_3$) and the hyperparameter on category probability $\nu_{\text{p}}$ as a free parameter as well. Here, instead we fit the model with $\nu_p = 2$, which was determined based on our experimental design. On average, this model provided a worse fit than the Wilson et al. (2013) model presented in the main text ($\Delta\text{LML}_{\text{covert}} = 28.79 \pm 9.91$; overt task: $\Delta\text{LML}_{\text{overt}} = 4.18 \pm 2.94$).

### 3 Model comparison with AIC

To ensure the robustness of our model comparison results, in addition to using the log marginal likelihood as a measure of goodness of fit, we calculated the Akaike information criterion (AIC) [4]. Unlike the log marginal likelihood, AIC uses a point estimate and penalizes for complexity by adding a correction for the number of parameters $k$: $AIC = 2k - 2\ln(\hat{L})$, where $\hat{L}$ is the maximum log likelihood of the dataset. Like the log marginal likelihood, AIC is best interpreted as a relative score. The model comparison results using relative AIC scores (relative to the winning model) are shown in Fig S1. From the plot we see that our results do not change using a different metric (compare with Fig 3 in the main text). Furthermore, the ranks for all models do not change for either task when comparing -0.5 AIC and LML scores ($\rho = 1.0$, $p < 0.0001$). Note that for historical reason the AIC scores have an additional factor of two.

### 4 Recovery analysis

#### 4.1 Model recovery

We performed a model recovery analysis to validate our model-fitting pipeline and ensure that models were identifiable [5]. For this analysis, we generated ten synthetic datasets from each model, observer, and task (1,980 datasets). Parameters for each simulated dataset were determined by sampling with replacement from the posterior over model parameters. We fitted these datasets with all models (17,820 fits), and for each pair of generating and fitting models we calculated the proportion of times each model fit the data best (i.e., had the greatest LML score), producing the confusion matrix in Fig S2. First, the fact that the confusion matrix is mostly diagonal means that most datasets were best fit by their true generating model, suggesting a generally successful recovery.

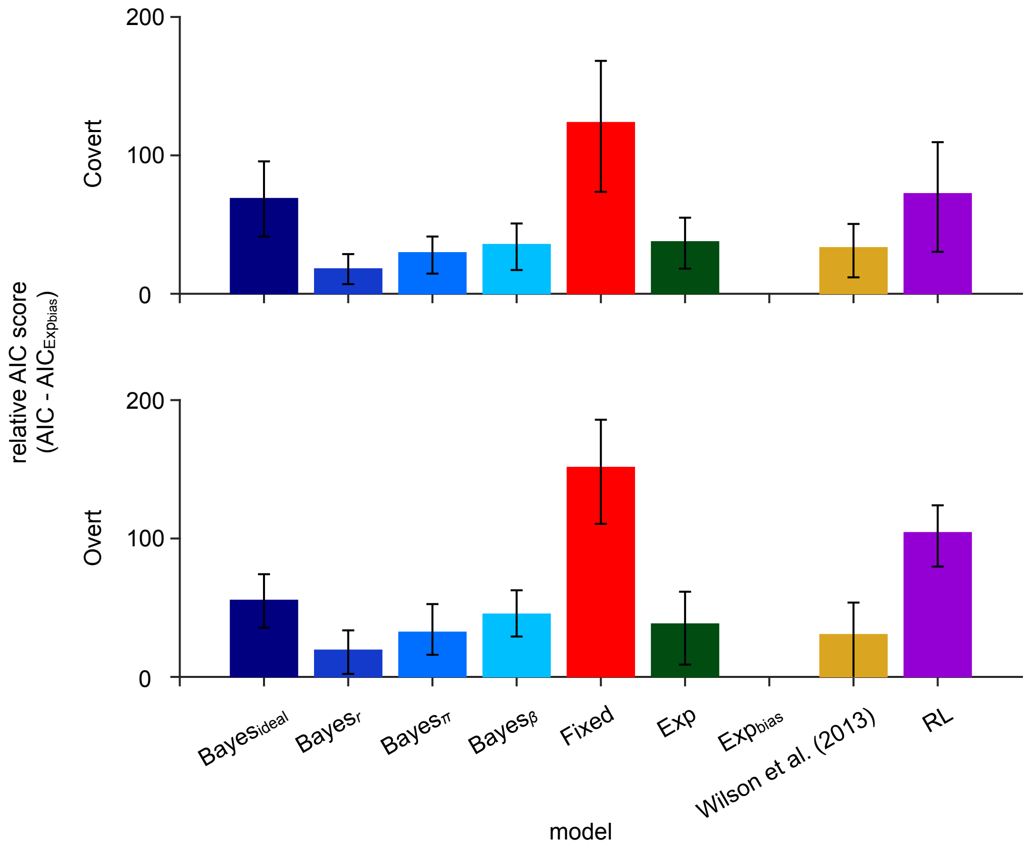

**Figure S1. Model comparison with AIC scores.** AIC scores relative to the $\text{Exp}_{\text{bias}}$ model are shown for the covert (top) and overt (bottom) tasks. Higher scores indicate a worse fit. Error bars: 95% C.I.

Across both tasks, we found that the true generating model was the best-fitting model for 70.1%±9.0% of simulated datasets (covert: 66.06%±11.36%; overt: 74.0% ± 8.6%; mean and SEM across models). For most simulated datasets, the true generating model was recovered for all models except the Exp model (see diagonal in Fig S2), which was best fit by the Wilson et al. (2013) model. However, this does not affect the results as the Wilson et al. (2013) model was not the best-fitting model across observers. Additionally, in the covert-criterion task (Fig S2B) the RL model simulations were best fit by the $\text{Exp}_{\text{bias}}$ model. This is potentially due to the fact that observers exhibited a greater amount of conservatism in the covert task. Increased conservatism results in smaller, smoother changes of criterion, which is consistent with what we observed in the RL model (see the bottom right panel in Fig 2D in the main text), so that data from the RL model are also well fit by the $\text{Exp}_{\text{bias}}$ model. However, these models were clearly distinguishable in the overt task (Fig S2C), which allows us to rule out the RL model. These results again provide support for the use of tasks, such as our overt task, that allow the researcher to better distinguish between computational models.

##### 4.2 Parameter recovery

To determine whether our parameter estimation procedure was biased, we analyzed the parameter recovery performance for the winning model ($\text{Exp}_{\text{bias}}$). Specifically, for each observer we created ten synthetic datasets by sampling from the posterior over model parameters and simulating the experiment with the same experimental parameters as the observer experienced. Each synthetic dataset was then fit to the $\text{Exp}_{\text{bias}}$ model and the best fitting parameters (MAP estimates) were estimated. For each parameter and task, we conducted a paired-samples t-test comparing the average best fitting parameters to the average generating parameters. We did not find a statistically significant difference between the fitted and generating $\alpha_{\text{Exp}}$ and $w$ parameters for either task: $\alpha_{\text{Exp}}$ (covert: $t(10) = 1.14$, $p = 0.28$; overt: $t(10) = 1.01$, $p = 0.34$) and $w$ (covert: $t(10) = -2.00$, $p = 0.07$; overt: $t(10) = 0.46$, $p = 0.66$), suggesting good parameter recovery. While there was no significant difference between the fitted and generating noise parameter ($\sigma_{\text{v}}$) in the covert task ($t(10) = 2.10$, $p = 0.06$), we found a significant difference in the noise parameter ($\sigma_{\text{a}}$) in the overt task ($t(10) = -16.53$, $p = 1.37e-08$). This difference remained significant after correcting for multiple comparisons using the Bonferroni cutoff of $p = 0.0083$. This result suggests that $\sigma_{\text{a}}$ was overestimated on average.

#### 5 Measurement task

##### 5.1 Procedure

During the ‘measurement’ session, observers completed a two-interval forced-choice, orientation-discrimination task in which two black ellipses were presented sequentially on a mid-gray background and the observer reported the

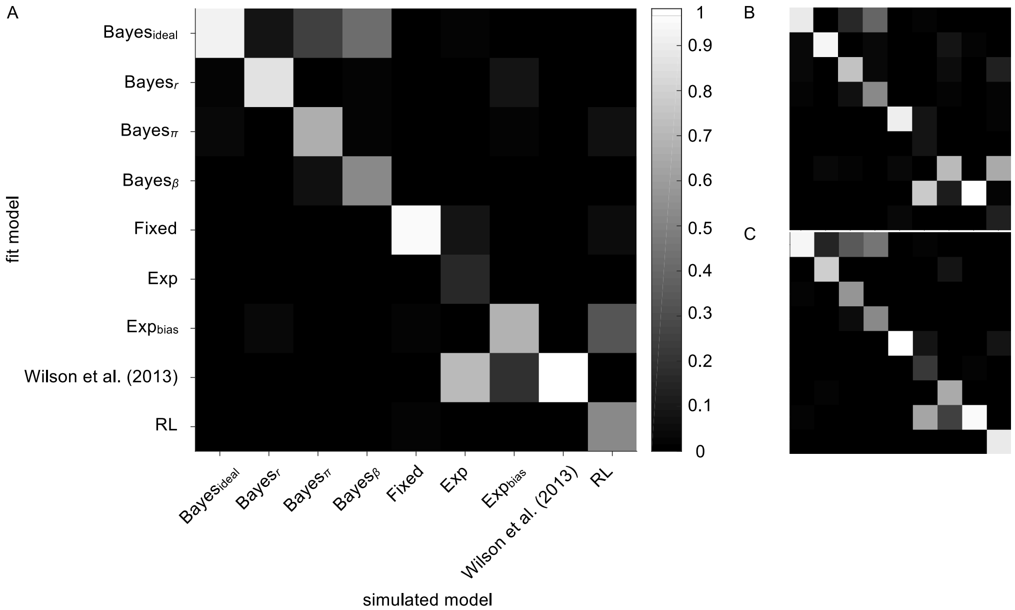

**Figure S2. Model recovery.** For each model, observer, and task, 10 sets of parameters were sampled from the model posterior and used to generate synthetic data (1,980 total simulations). The synthetic datasets were then fit to each model (17,820 fits) and the goodness of fit was judged by computing the LML. The proportion of "wins" (i.e., the number of times the simulated model outperformed the alternative models) is indicated by brightness. Model recovery performance is shown across both tasks (A), the covert task only (B), and the overt task only (C).

interval containing the more clockwise ellipse (Fig S3A). This allowed us to measure the observer's sensory uncertainty.

#### 5.2 Analysis

A cumulative normal distribution was fit to the orientation-discrimination data (probability of choosing interval one as a function of the orientation difference between the first and second ellipse) using a maximum-likelihood criterion with parameters $\mu$, $\sigma$, and $\lambda$ (the mean, SD, and lapse rate). We define threshold as the underlying measurement SD $\sigma_{\text{v}}$ (correcting for the 2IFC task by dividing by $\sqrt{2}$).

#### 5.3 Results

Fig S3B shows a representative psychometric function for one observer. The average threshold across observers was $\sigma_{\text{v}} = 6.71° \pm 1.23°$.

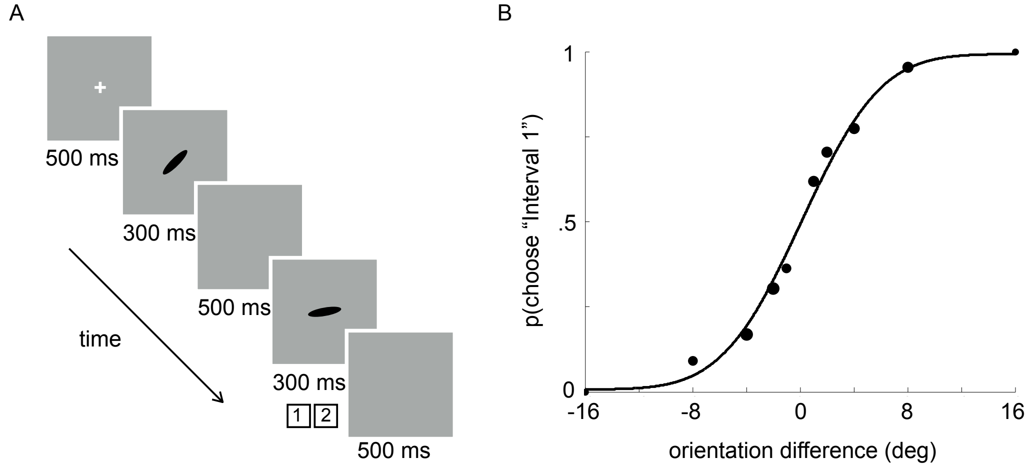

**Figure S3. Measurement task.** A: Trial sequence. Two ellipses were presented sequentially on a mid-gray background. The observer reported the interval containing the more clockwise ellipse. Feedback was provided. B: The best fitting psychometric function for one observer. The area of each data point is proportional to the number of trials.

### 6 Category training

#### 6.1 Procedure

Category training was completed prior to the covert- and overt-criterion tasks, so observers could learn the category distributions. It was important not to confound category learning with probability learning. Training was identical to the covert-criterion task (Fig S4A). On each trial ($N_{\text{trials}} = 200$), a black ellipse was presented on a mid-gray background and observers reported the category to which it belonged. Category probability was equal during training and we provided correctness feedback. To determine how well observers learned the category distributions, observers estimated the mean orientation of each category at the end of the training block by rotating an ellipse to match the mean orientation (Fig S4B). Each category was estimated exactly once.

#### 6.2 Results

Observers' estimates of the category means are shown in Fig S4C as a function of the true mean. Data points represents each observer's estimate after each task for each category. There was a significant correlation between category estimates and the true category means (category A: $r = 0.82$, $p < 0.0001$; category B: $r = 0.97$, $p < 0.0001$), suggesting that participants learned the categories reasonably well. On average, estimates were repelled from the category boundary (average category A error of $11.3° \pm 6.3°$ and average category B error of $-8.0° \pm 2.6°$; mean and SEM across observers).

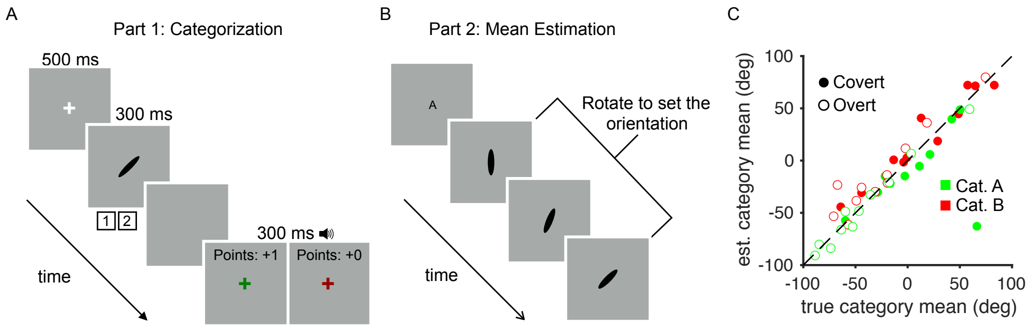

**Figure S4. Category training.** A: Trial sequence. After stimulus offset observers reported the category by key press and received feedback. B: Mean estimation task. After completing the training block, observers rotated an ellipse to estimate the category means. C: Estimation results. Observers' category-mean estimates are shown as a function of the true category mean for each category, observer and task.
